# The NHEJ machinery is translocated off DNA ends to enable resection

**DOI:** 10.1101/2025.05.13.653820

**Authors:** Pradeep Sathyanarayana, Yufan Wu, Johannes C. Walter, Joseph J. Loparo

## Abstract

Error-prone non-homologous end joining (NHEJ) and homologous recombination (HR) compete to repair DNA double strand breaks [1, 2, 3]. Given that NHEJ factors such as Ku and DNA-PKcs are rapidly recruited to DSBs and sterically occlude ends, they must be removed to enable 5^*′*^ DNA end resection, the commitment step of HR. While resection is well characterized, how the NHEJ machinery is displaced from DNA ends has remained poorly understood. Here, we performed single-molecule imaging in *Xenopus laevis* egg extracts to monitor the removal of NHEJ factors from DNA ends while simultaneously tracking the progress of DNA end resection. We find that MRN and CtIP are essential to evict the core NHEJ factor Ku from DNA ends. Surprisingly, while Ku removal and resection of the 5^*′*^ end are normally coordinated, these processes arise from distinct activities of the MRN complex and are differentially regulated by the action of cyclin dependent kinases and ATM. By directly imaging the position of Ku on DNA using smFRET, we found that Ku is translocated independently of DNA end resection in an ATP-dependent manner that requires both MRN and CtIP. In summary, our results provide a new mechanistic framework to understand how eviction of the NHEJ machinery is coordinated with DNA resection to initiate HR.

## Introduction

DNA double strand breaks (DSBs) are toxic lesions that must be properly repaired to avoid genome rearrangements [4]. DSB repair is carried out by two major pathways: non-homologous end joining (NHEJ) [5] and homologous recombination (HR) [6]. NHEJ is initiated by the binding of Ku and DNA-PKcs to DNA ends [7, 8, 9]. Subsequent recruitment of core and accessory NHEJ factors results in DNA end synapsis [10], processing [11] and ligation [12, 13]. As end processing can lead to insertions and deletions, NHEJ is often considered error-prone [5, 14]. In HR, 3^*′*^ single-stranded (ss) DNA overhangs are generated by nucleolytic degradation of the 5^*′*^ end in a process known as resection [15]. The resulting ssDNA is ultimately coated by the Rad51 recombinase which mediates strand-invasion into a homologous region in the sister-chromatid and primes error-free repair [3].

Resection is initiated by the Mre11-Rad50-Nbs1 (MRN) complex along with its essential co-factor CtIP [16, 17]. The MRN complex is composed of the Mre11 nuclease, the Rad50 ATPase which is an SMC family member and the structural cofactor Nbs1 (Nijmegen Breakage Syndrome) [18, 19]. Mre11 possesses 3^*′*^→ 5^*′*^ exo- as well as endonuclease activities [20], which are regulated by Rad50 and CtIP [20, 21]. Phosphorylation of CtIP by cyclin dependent kinase (CDK) stimulates MRN endonuclease activity, thereby restricting DNA resection to the S and G2 phases of cell cycle [22, 23]. CtIP is further phosphorylated by the DNA damage kinase ATM, which is also indispensable for resection [24]. Resection is believed to occur through a bidirectional model in which an initial endonucleolytic nick is made interior to the DSB. Subsequent short-range resection mediated by MRN occurs in the 3^*′*^ →5^*′*^ direction while long-range resection is carried out in the 5^*′*^ →3^*′*^ direction by the processive nucleases EXO1 and DNA2 [25, 26]. DNA end resection is a critical step in committing repair to HR, as it makes the ssDNA overhangs refractory to Ku loading, thereby inhibiting NHEJ [3, 26, 2].

The NHEJ components Ku and DNA-PKcs are rapidly recruited to breaks throughout the cell cycle due to their high abundance and affinity for DNA ends [27, 28, 29]. Ku encircles dsDNA and DNA-PKcs locks it in place near the DNA end [30] which protects the ends from processing and resection [31]. In the S and G2 phases, where both NHEJ and HR pathways are active [23, 32], removal of the NHEJ complex is essential for repair to proceed via the error-free HR pathway. While resection is implicated in Ku removal [33, 34, 35], the mechanism by which nucleolytic degradation is coordinated with the eviction of the NHEJ machinery has remained unclear.

By carrying out single-molecule imaging experiments that probe both NHEJ complex removal and 5^*′*^ DNA end resection in *Xenopus* egg extracts, we reveal the mechanism by which the NHEJ machinery is removed from ends to facilitate DNA resection. We find that Ku removal precedes 5^*′*^ end resection by a short time interval. We further demonstrate that MRN mediates eviction of the NHEJ machinery through an active translocation mechanism that is independent of its known resection activity. These distinct functions of the MRN complex are differentially regulated by CtIP phosphorylation and are mediated through interactions with Rad50. While translocation activity is activated by CDK-dependent phosphorylation of CtIP, resection only occurs upon subsequent ATM phosphorylation. Collectively, our observations report on a previously unknown function of the MRN complex that is essential to its role in the initiation of DNA end resection and HR.

## Results

### Single-molecule eviction of the NHEJ complex from DNA ends

To study the mechanism of eviction of the NHEJ machinery from DNA ends, we designed a single-molecule assay using *Xenopus laevis* egg extracts. A 1.5kb dsDNA containing a ATTO647 fluorophore 5 nt from the 5^*′*^ end was tethered to the surface in a microchannel. The NHEJ machinery was loaded onto the DNA ends within the microchannel by introducing a high-speed supernatant of egg extracts (HSS) [36] that mimics the G1 phase of the cell cycle. The endogenous Ku in G1 extract was immunodepleted and replaced with fluorescently labeled Ku (Halo[JF549]-Ku) [37] (Fig. 1a). We previously demonstrated that the G1 extract supports both synapsis of broken DNA ends and end processing, resulting in robust NHEJ [10, 11]. Ku was rapidly recruited to DNA molecules (∼ 1 min) and single-molecule photobleaching analysis of the JF549 spots indicated that most DNA molecules contained a single copy of Ku [37] (Fig. 1b). To study the HR mediated eviction of the NHEJ complexes, we introduced an S-phase extract (NPE; nucleoplasmic extract) into the microchannel. Removal of the NHEJ machinery was observed as a step loss of labelled Ku fluorescence from DNA ends (Fig. 1c). Survival probability curves showed that the NHEJ machinery was rapidly removed in the S-phase extract while being stably associated in the G1 extract, consistent with the fact that HR is restricted to S/G2 phases of cell cycle [3] (Fig. 1d).

**Figure 1:**
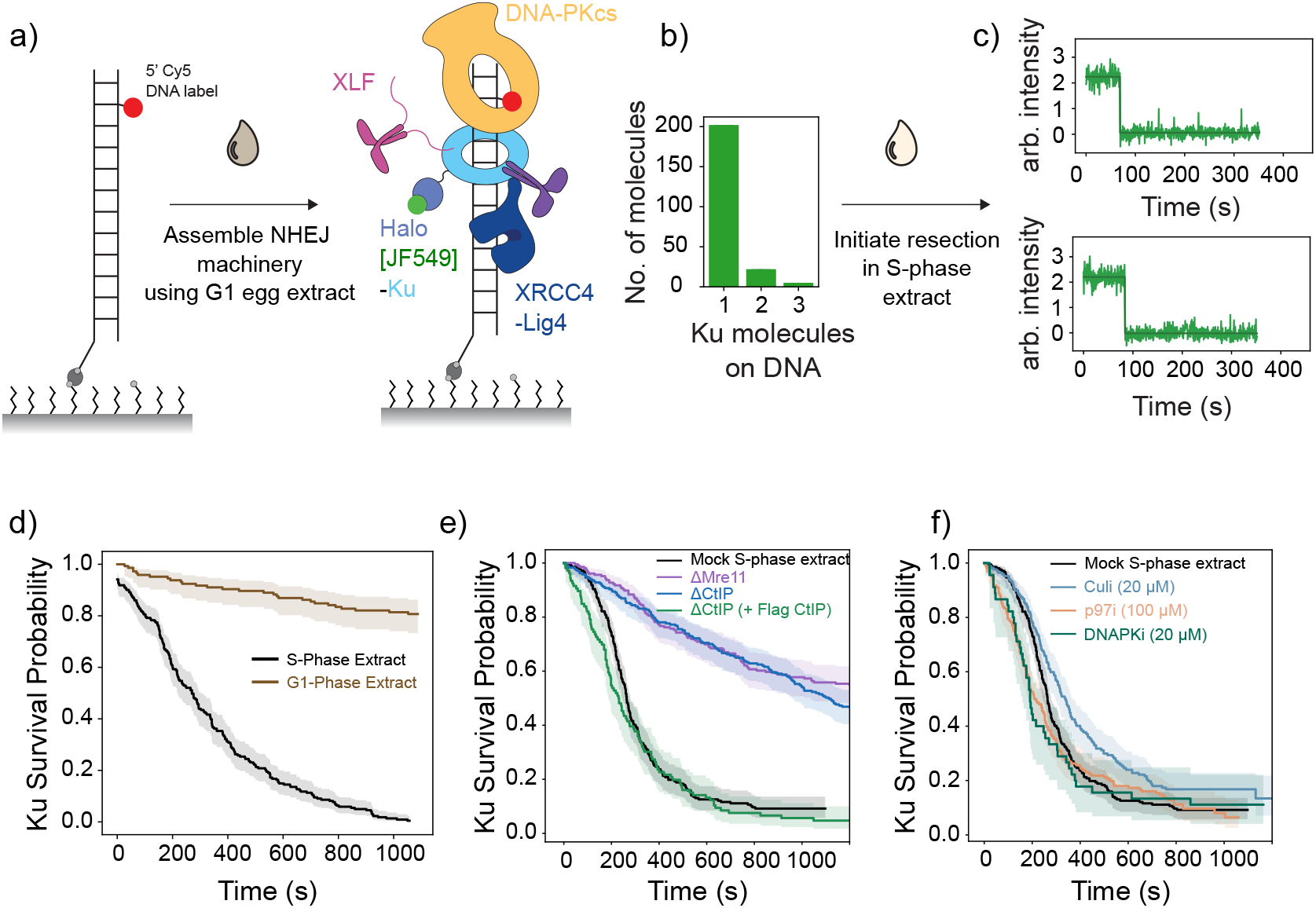
Single-molecule assay to study NHEJ complex eviction. a) Schematic representation of the single-molecule experiment. A 1.5kb DNA tethered to glass surface is loaded with the NHEJ complex by flowing G1 phase egg extracts containing fluorescently labeled Ku (JF549) into the flow cell. B) Histogram of Ku stoichiometry assessed by photobleaching analysis showing that most DNA molecules contain a single molecule of Ku (see SI, methods) C) Fluorescent intensity traces of labelled Ku after introducing S-phase egg extracts into the flow cell. The step-change in the trace represents the loss of a single-molecule of Ku. D) Survival probability of Ku on DNA ends in G1 and S-phase egg extracts (see methods). Ku molecules are rapidly removed in the S-phase extract. E) Ku survival probability in S-phase extracts supplemented with indicated inhibitors. Concentrations used are indicated within brackets. F) Ku survival probability in indicated S-phase extracts. Recombinant Flag-CtIP was added to a final concentration of 30 nM. Shaded regions in all survival graphs represent 95% confidence interval.

Several pathways have been implicated in removing the NHEJ machinery from DNA ends, including MRN-initiated DNA resection and posttranslational modifications of Ku such as ubiqutination and phosphorylation [34, 35, 38, 39]. Since egg extracts support all the aforementioned processes, we asked if and how these pathways contribute to NHEJ machinery removal in S-phase extract. Individual immunodepletions of MRN complex (using Mre11 antibody), and CtIP resulted in a drastic reduction in Ku removal with >50% molecules being retained on ends after long times (>1200 s) (Fig. 1e). While addition of recombinant MRN complex failed to rescue Ku removal in Mre11 depleted extracts, Ku eviction was restored to normal levels by adding back recombinant CtIP to immunodepleted S-phase extracts (Fig. 1e and Extended data Fig. 1), demonstrating that the defect in Ku removal was due to the absence of CtIP. The contribution of phosphorylation and ubiquitination towards Ku removal was examined using specific small molecule inhibitors against DNA-PKcs (NU7441; kinase inhibitor), Cullin ring ligase (MLN4924) and p97 (NMS873) (Fig. 1f). While DNA-PKcs and p97 inhibition had little effect on NHEJ complex eviction, Cullin inhibition showed a slight defect (Table.1) in the kinetics of NHEJ complex removal but eventually resulted in ∼ 0% of molecules being lost. Together, the results indicate that NHEJ is evicted from ends in a resection dependent manner, with limited contribution from other processes such as ubiqutination or phosphorylation.

**Table 1:**
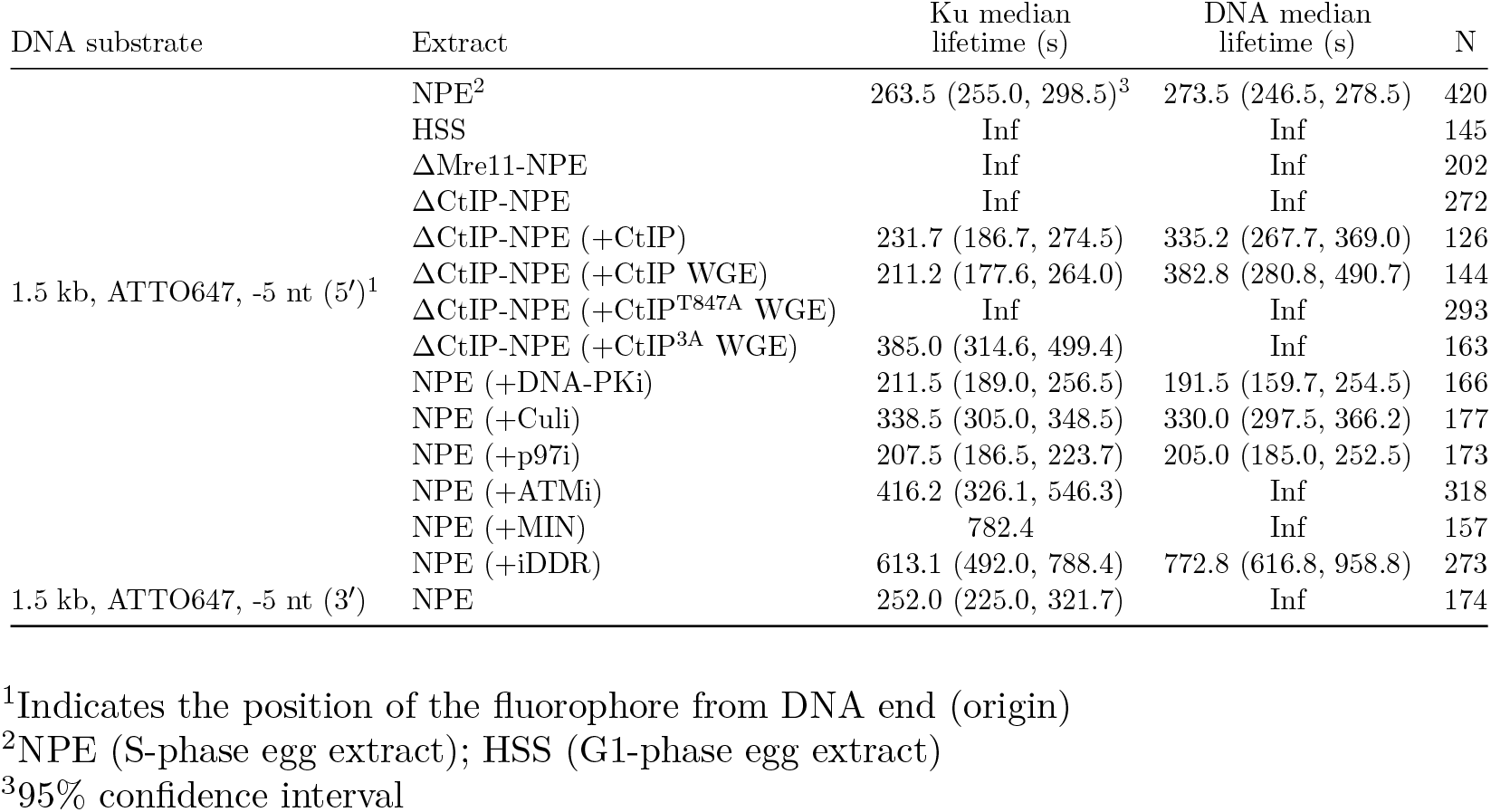
Quantification of Ku removal and 5^*′*^ end resection.

### Resection and NHEJ complex eviction are coordinated processes

To understand the coordination between DNA resection and NHEJ complex removal, we monitored the loss of the fluorescently labeled 5^*′*^ DNA strand and Halo [JF549]-Ku in S-phase extracts. On most DNA molecules Ku eviction was soon followed by resection of the 5^*′*^ strand (Fig. 2a). Quantification of the lag time between Ku removal and resection across all DNA molecules showed an average time of -8.22 s (−11.77, -4.94; 95% CI) (Fig. 2b). We also observed a fraction of molecules (14.82 %) where both processes occured simultaneously. Collectively, these results demonstrate that NHEJ complex removal is critical for resection to proceed, consistent with prior observations that Ku blocks resection and its removal is essential for resection to proceed [40].

**Figure 2:**
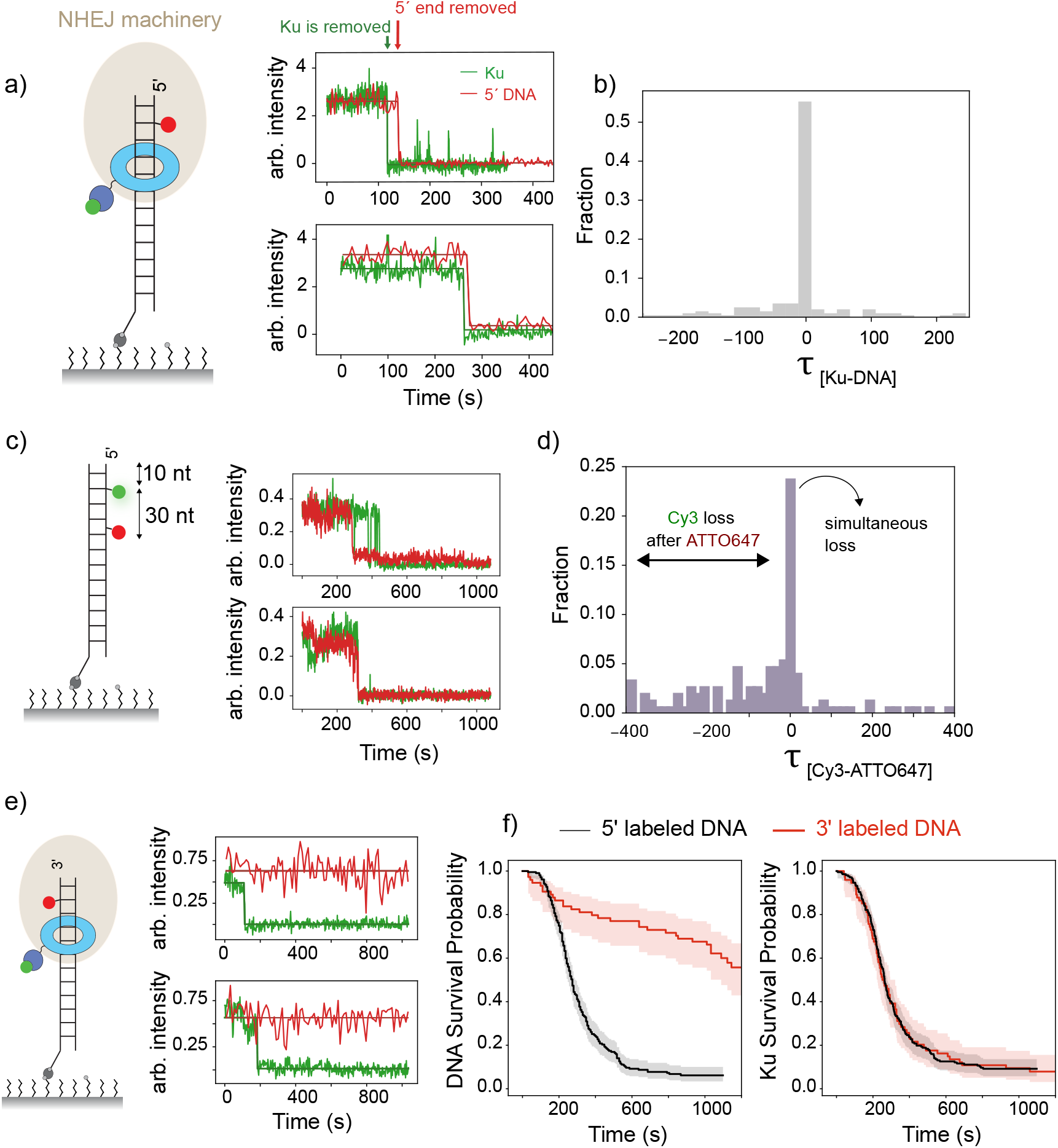
NHEJ complex removal and 5^′^ end resection are coordinated processes. Schematic representation of the NHEJ complex containing Halo[JF549]-Ku loaded on DNA with 5^*′*^ ATTO-647 label. On the right, two representative graphs containing fluorescent intensity traces of labelled Ku (green) and 5^*′*^ DNA end (red). Resection of 5^*′*^ end follows shortly after loss of Ku from DNA. b) Histogram of lag-time between Ku and 5^*′*^ end loss (see Table. 1) in S-phase egg extract c) Schematic of DNA labelled with Cy3 and ATTO647 fluorophores at indicated positions. Representative graphs showing Cy3 (green) and ATTO647 (red) intensity traces of the substrate (left) in S-phase egg extract. d) Histogram of lag-time between Cy3 and ATTO647 loss from Fig. 2C. e) Schematic of DNA substrate labelled with ATTO-647 on 3^*′*^ end. Representative graphs containing intensity traces of Ku (green) and 3^*′*^ end (red) in S-phase extract. f) DNA (left) and Ku (right) survival probability from Fig. 2E. Shaded represent 95% confidence interval.

Next, we asked if DNA end resection requires the exonuclease activity of Mre11 or if its endonuclease activity is sufficient. To investigate this, we engineered a 1.5 kb DNA substrate with Cy3 and ATTO647 fluorophores at locations 10 and 40 nucleotides from the end, respectively (Fig. 2c). As described before (Fig. 1a) the DNA substrates were loaded with the NHEJ machinery before addition of the S-phase extract and imaging was carried out by simultaneous excitation of the fluorophores.

Frequently, both dyes were lost simultaneously, indicating that relatively long oligos (∼ 30 nt) were released as resection products (Fig. 2c, d). For most other molecules, the internal ATTO647 (−40 nt) was lost before Cy3 (−10 nt). Taken together, this points towards resection likely arising from multiple endonucleolytic nicks near the 5^*′*^ end, initiating internally and extending towards the break. Alternatively, simultaneous loss of the two fluorophores could result if endonucleolytic cleavage on both strands released a dsDNA fragment containing the NHEJ complex, as previously reported [33]. To directly test for the dsDNA cleavage mechanism, we moved the ATTO647 flurophore to the 3^*′*^ strand (−5 nt) and monitored Ku removal and resection. While the kinetics of Ku removal remained unaffected (Fig. 2e), the 3^*′*^ fluorophore exhibited far greater stability as compared to when the DNAs were labeled on the 5^*′*^ end (Fig. 2f). To further verify that a 3^*′*^ overhang is retained upon resection of the 5^*′*^ end, we washed the flow cell with 2% SDS to remove all proteins and flowed in Cy3 and Cy5 labeled oligonucleotides that were complementary to different segments (Cy3oligos-body; Cy5oligo-end) of the expected 3^*′*^overhang. The colocalization of the two oligos indicated that the ssDNA overhang was retained on the surface (Extended data Fig. 2). In conclusion, our data shows that the short-patch resection is achieved by a combination of multiple MRN-mediated nicks near the 5^*′*^ end.

### NHEJ eviction and DNA resection are independently regulated

DNA resection initiated by the MRN complex requires the phosphorylation of CtIP by CDK and ATM [16, 24] (Fig. 3a). We first asked how CDK phosphorylation affects the removal of Ku and end resection. We generated a phosphoablating mutant of the CDK site on CtIP^T847A^ (Fig. 3b) and expressed the full-length protein using an in-vitro transcription and translation system (Wheat germ extract; WGE). After verifying expression (Extended data Fig. 1b), ΔCtIP S-phase extract was supplemented with CtIP^T847A^ or WT (10% v/v WGE) and assessed for NHEJ complex removal and resection as in Fig. 1a. Unlike the WT, CtIP^T847A^ (Fig. 3b) failed to rescue Ku removal (Fig. 3c; left) or resection (Fig. 3c; right). As CtIP is further phosphorylated by ATM in response to DNA damage, we used a selective ATM inhibitor (KU-55933) to determine the role of ATM signaling on Ku removal and DNA resection. While ATMi significantly inhibited resection (Fig. 3d; right), Ku was still removed from DNA ends, albeit with slightly slower kinetics (t_1/2_ = 451 s) (Fig. 3d; right). To determine if this defect in resection was due to loss of CtIP phosphorylation, we introduced three phosophoablating mutations into conserved SQ/TQ sites in the C-terminus of *X. laevis* CtIP (CtIP^3A^ (664A/679A/859A); residue numbers correspond to human CtIP) [24]. ΔCtIP S-phase extract supplemented with the CtIP^3A^ mutant was significantly defective in 5^*′*^ end resection (Fig. 3d; left) while retaining the ability to remove Ku from DNA ends, phenocopying the ATMi results (Fig. 3d; right, Table. 1).

**Figure 3:**
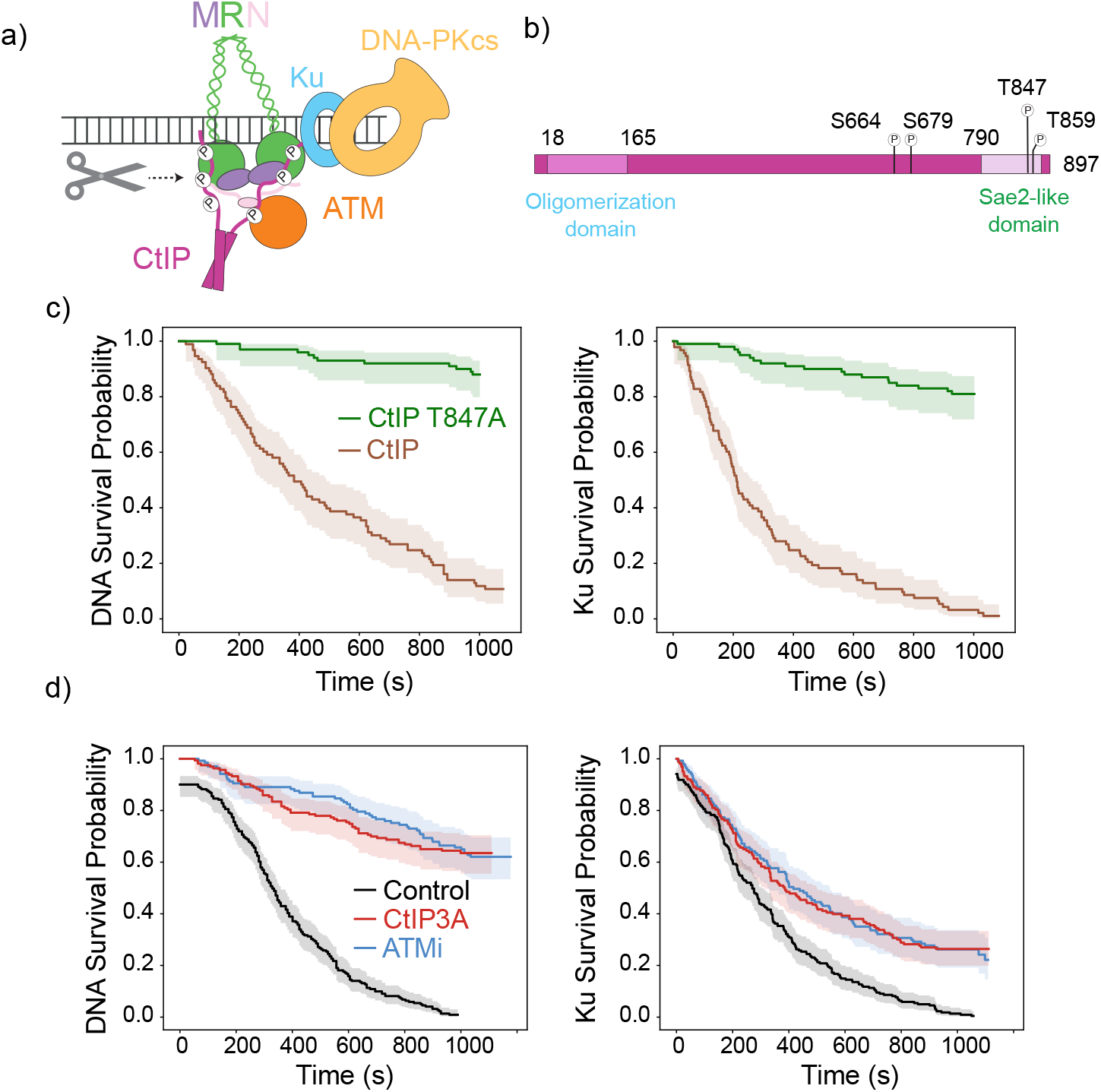
CtIP phosphorylation by CDK and ATM regulates Ku removal and 5^′^ end resection respectively. a) Schematic highlighting the key components of the resection machinery on DNA ends bound by the NHEJ complex. b) Schematic representation of the various domains of human CtIP. S664, S679 and T859 are ATM sites and T847 is phosphorylated by CDK. c) Ku (left) and 5^*′*^ DNA (right) survival probability in ΔCtIP S-phase extracts supplemented with CtIP (green) or CtIP^T847A^ (brown). d) Ku (left) and 5^*′*^ DNA (right) survival probability of S-phase extract supplemented with ATM inhibitor (KU-55933) at a concentration of 20 µM or ΔCtIP S-phase extract supplemented with ATM site mutant, CtIP^3A^. CtIP^3A^ mutant phenocopies the ATMi inhibition. CtIP, CtIP^T847A^ and CtIP^3A^ mutants used in the experiment were expressed in WGE (see methods, SI). Shaded regions in all survival graphs represent 95% confidence interval.

While current models stipulate that Ku removal is a consequence of 5^*′*^ end resection [35, 26, 2], our results demonstrate that Ku removal and 5^*′*^ end resection involve distinct activities. CtIP phosphorylation by CDK is essential for removal of the NHEJ machinery and resection while ATM phosphorylation is specifically necessary to initiate resection. Since CtIP lacks nuclease activity [17], we hypothesized that CtIP exerts its effect via interaction with the MRN complex.

### Phosphorylated CtIP-Rad50 interaction is essential for NHEJ complex removal and resection

To understand how phosphorylated CtIP interacts with the MRN complex, we folded the human MR complex, pCtIP and dsDNA (60 bp) using AlphaFold multimer V 3.0 [41]. In the absence of phosphorylated CtIP, MR adopted the inactive cutting state [42] (4 out of 5 models) in which the Mre11 dimer is positioned below the Rad50 globular domains (Fig. 4a and Extended data Fig. 3). However, when folded with CtIP phosphorylated on CDK (T847) and ATM (S664, S679 and T859) sites, Mre11 was repositioned behind the Rad50 globular domains resulting in the catalytic H129 being directed towards the DNA (Fig. 4a, right). This conformation is reminiscent of the cutting state of the *E. coli* MR complex [42]. The coiled coils of Rad50 are constricted and collapse around dsDNA compared to the more open conformation in the model without CtIP. Notably, the model predicts a conserved interface between the Rad50 globular domain and the C-terminus of CtIP with the CDK site (T847) and the crucial ATM site (T859), both packed against a positively charged pocket in the N-terminal Rad50 ATPase domain (SI, Fig. 4b). K6 and R79 of Rad50 make electrostatic interactions with T859 and T847 of CtIP, respectively. Interestingly, Rad50K6E is a rad50s allele that results in embryonic lethality in mice, highlighting that the predicted AF3 interface is crucial for HR [43]. To examine the interaction between CtIP and Rad50, we carried out pull-down experiments with purified xMRN complex and CtIP expressed using WGE. CtIP^T847A^ and CtIP^3A^ were both significantly compromised in their ability to bind to the MRN complex (Fig. 4c). The small amount of CtIP that was pulled down was dependent on phosphorylation, and is possibly mediated via interaction with the Nbs1 subunit [20, 24]. To further validate this interface, we used peptides derived from telomeric proteins that prevent ATM signaling by binding to the Rad50 subunit of the MRN/X complex and disrupting its interaction with Sae2. The yeast Rif2 protein harbors a short amino acid motif termed MIN (MIN; **M**RN **IN**hibitor) [44] that disables MRX activity, and an analogous motif (called iDDR) [45] has been recently described in the human TERF2. Furthermore, AlphaFold models of the MIN peptides with Rad50 showed that they mapped to the same positively charged *β*-sheet region in the N-terminus of Rad50 that made contacts with CDK and ATM phosphorylation sites on CtIP (Extended data Fig. 4b, c).

**Figure 4:**
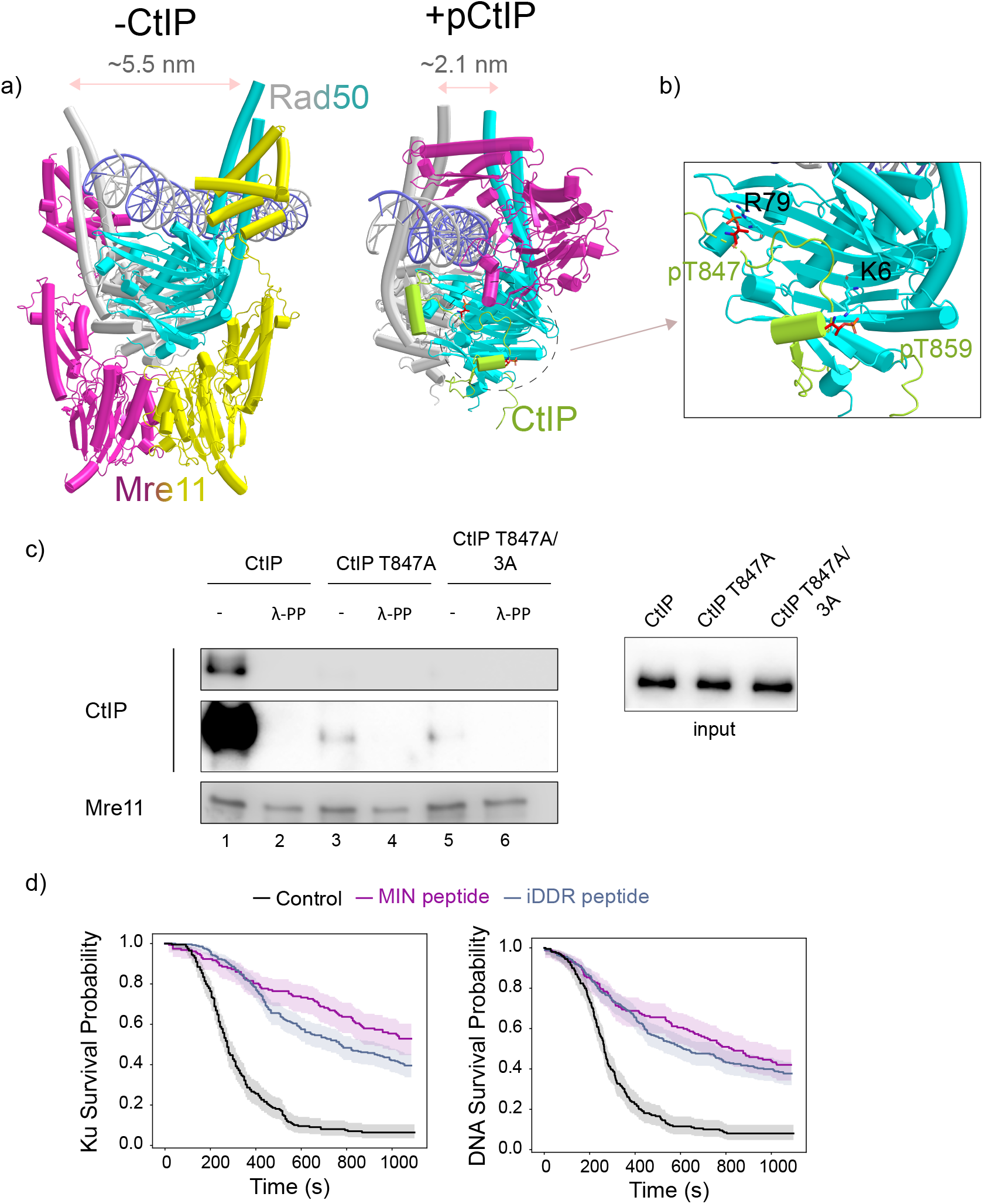
CtIP phosphorylation modulates Ku removal and resection via interaction with Rad50. a) AlphaFold 3 model of Mre11-Rad50 (left; inactive state) and Mre11-Rad50-pCtIP (right; active state) folded with 60 bp dsDNA. Only residues having PLDDT values >50 are displayed. In the active state, only a single protomer of Mre11 is displayed for clarity. The presence of phosphorylated CtIP favors the active cutting state. Highlighted in dashed circle is the conserved binding interface between CtIP and Rad50. Zoomed in view highlighting the key residues that are involved in interaction between CtIP and Rad50. c) Protein interaction assay with purified xMRN complex and WGE expressing CtIP or mutants. Interacting proteins were pulled down and detected by Western blotting using anti-CtIP or anti-Mre11 antibodies. d) Ku (left) and 5^*′*^ DNA (right) survival probability in S-phase extracts that are supplemented with MIN (250 µM) and iDDR peptide (300 µM) (see methods and SI). Shaded regions represent 95% confidence interval.

We hypothesized that if the interaction between Rad50 and CtIP was crucial for NHEJ eviction, S-phase extracts spiked with peptides derived from the MIN or iDDR sequences would inhibit Ku removal. We found that not only was NHEJ removal and resection inhibited in the presence of *X. laevis* TERF2 iDDR, but the heterologous *S. cerevisiae* MIN peptide also exhibited a drastic inhibitory effect (Fig. 4d). In contrast, scrambled iDDR peptide sequence had little effect on Ku removal and end resection kinetics (Extended data Fig. 5). Together these results demonstrate that phosphorylation of CtIP by CDK and ATM at distinct sites modulates its interaction with specific residues in the Rad50 N-terminus which in turn regulates NHEJ eviction and DNA resection independently.

### NHEJ complex is translocated off DNA ends prior to resection

Our results show that Ku removal can still occur in the absence of DNA resection. Since Rad50 belongs to the SMC family of proteins, which includes known loop extruders such as cohesin, Smc5/6 and condensin [46, 47, 48], we postulated that the NHEJ machinery is mechanochemically removed from DNA ends. To test if motor activity was involved in NHEJ complex removal, we designed a single-molecule FRET assay using donor-labeled Ku and acceptor-labeled DNA (Fig. 5a). If motor activity pushes Ku off the DNA end (rather than opening of the Ku clamp and lateral dissociation), we expected to see an increase in FRET between Ku and a fluorophore near the DNA end. Ku70 was site-specifically labelled with a Cy3 dye using a ybbr tag inserted in a loop region proximal to DNA so that FRET would be observed upon Ku binding near a ATTO647 fluorophore at the 5^*′*^ end of the DNA (Fig. 5a, b). We further verified that incorporation of the ybbr tag and subsequent fluorescent labeling did not compromise the activity of Ku in end-joining (Extended data Fig.6a and b). NHEJ complex was loaded using ΔKu-G1 extract supplemented with Ku80/Ku70-Cy3 and bound molecules showed an initial low-FRET state corresponding to a FRET efficiency of ∼ 0.3. Upon addition of S-phase extract, we observed a sharp increase in FRET efficiency to ∼ 0.9, which was followed by the sudden loss of the donor signal due to the removal of Ku from DNA ends (Fig. 5c and Extended data Fig. 7). While the Ku lifetime on DNA was ∼ 263 s (see Fig. 1D, Table. 1), the transition from low to high-FRET occurred over a shorter period of ∼ 60 s. Shortly after Ku was lost, resection was typically observed as seen in the stepwise loss of the acceptor signal (Fig. 5c, bottom panel). To further verify that the change in FRET was indeed due to the motion of Ku towards the end, we shifted the position of the acceptor fluorophore, from 5 nt from the end on the 5^*′*^ strand to -25 nt. In this configuration, FRET should decrease as Ku translocates towards the end. As expected, the FRET efficiency upon Ku binding was ∼ 0.4 and showed a steady decrease with eventual loss of Ku occuring at FRET efficiency of ∼ 0.05, a reversal in the trend seen in Fig. 5a (Extended data Fig. 8). When ATM inhibitor was added to S-phase extract to block resection, Ku was still translocated off the DNA ends (Fig. 5d), but exhibited increased dynamics prior to its removal (Extended data Fig. 9). Ku translocation and resection processes were both impaired in ΔCtIP S-phase extracts supplemented with the CDK phosphoablating mutant CtIP^T847A^ (Extended data Fig. 10) highlighting their inability to stimulate Rad50 motor activity.

**Figure 5:**
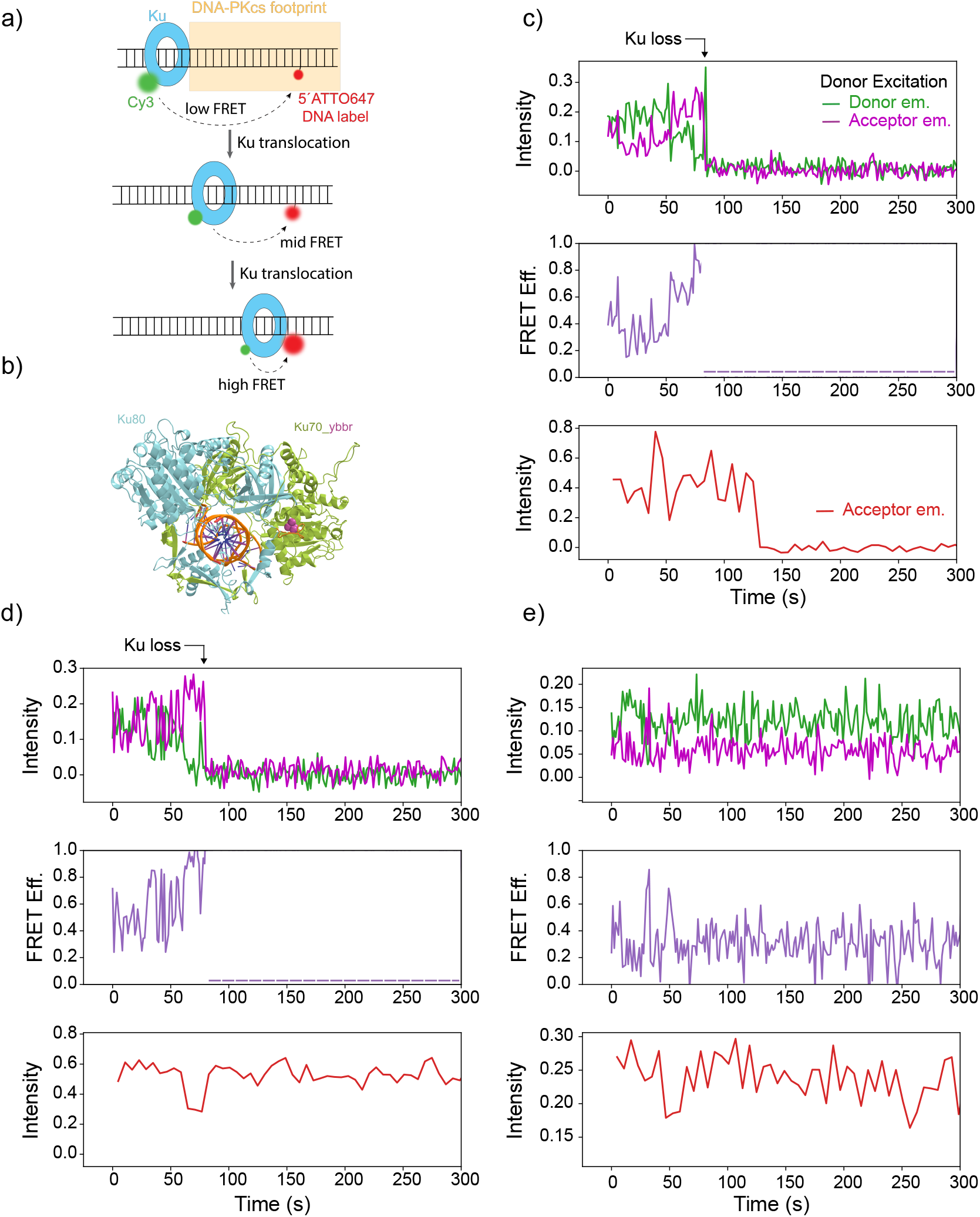
Ku is mechanochemically translocated off DNA ends. a) Schematic representation of smFRET assay to monitor Ku translocation. Ku translocation towards the DNA end will result in progressive increase in FRET efficiency. b) AlphaFold 2 model of Ku80/Ku70-ybbr (see methods) aligned to PDB ID:1EJY. Maroon sphere indicates position of fluorophore. c, d and e) Intensity traces from donor (green; Ku) and acceptor emission (maroon) upon donor excitation (*top*) and, acceptor emission (red; 5^*′*^ DNA end) upon acceptor excitation (*bottom*). The middle panel highlights the calculated FRET efficiency trace. c) Representative trajectory for experiment described in Fig. 5a in S-phase extract. FRET efficiency trace shows an increase from low- (E ∼ 0.3) to high-FRET state (E ∼ 0.9) after which Ku is lost, and immediately followed by loss of 5^*′*^ end. Representative trajectory in S-phase extract supplemented with, d) 20 µM ATM inhibitor and, E) slowly hydrolysable ATP analog (ATP*γ*S; 3 mM)

Since ATPase activity is essential for motor function of the SMC family proteins, we posited that spiking the S-phase extract with the slowly hydrolysable ATP analog, ATP*γ*S would result in inhibition of Ku removal. Consistent with this idea, Ku was locked in the low-FRET state and translocation was inhibited (Fig. 5e and Extended Fig. 11). ATP hydrolysis by Rad50 is essential for conformational rearrangement of Mre11 from the inactive state to access DNA ends to carry out the endonucleolytic incision (Fig. 4a). As a result, the ATP*γ*S addition to S-phase extracts also resulted in substantial inhibition of 5^*′*^ end resection. In summary, these results demonstrate that the NHEJ complex is actively pushed off DNA ends likely by Rad50 motor activity which is in turn regulated by interaction with phosphorylated CtIP.

### Multiple attempts are required for successful eviction of Ku

In the FRET efficiency traces (Fig. 5C and D), we observed several short-lived transition events at intermediate FRET values. When the experiment was carried out at much faster (24-96x) imaging conditions (10-50 ms exposure time), oscillatory motions of Ku became more evident; FRET trajectories transitioned from a low-FRET state (E = ∼ 0.25) to an intermediate state (E= ∼ 0.55) and back (Fig. 6a).

**Figure 6:**
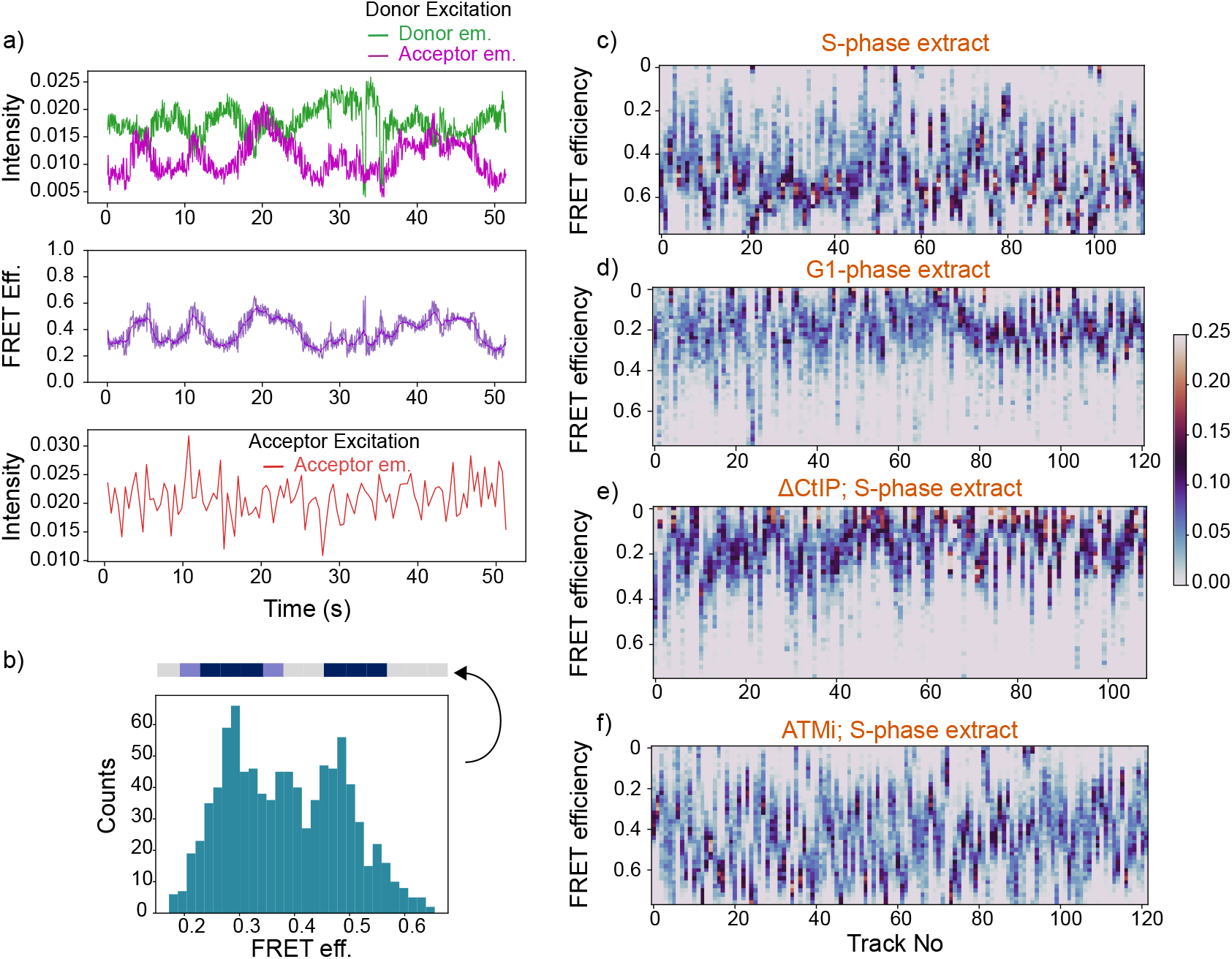
Multiple translocation attempts are required for the successful removal of Ku. Same as Fig. 5c but image capture was carried out with integration time of 25 ms between frames (see methods, SI). Traces from a representative molecule showing oscillating donor intensity (top panel) indicating the underlying Ku motion. FRET efficiency trace shows recurrent transitions from low-FRET state to intermediate-FRET state (middle panel). Denoised and smoothed FRET efficiency trace (see methods) is shown in continuous violet line. b) Histogram of the FRET efficiency from Fig. 6a. 1D schematic representation of this histogram is shown above the graph. c, d, e and f) 1D histogram of FRET efficiency from multiple molecules were concatenated (column wise) to generate the image. Color bar indicating the probability density from the original histogram is shown on the right. Each image is generated by analyzing traces obtained from different extract conditions as indicated.

The number of oscillatory cycles between the two states was variable (Extended Fig. 12) and the transition between the two was a smooth progression with one cycle lasting ∼ 5s (see discussion). A histogram of the FRET efficiency from the trace in Fig. 6a shows the presence of two peaks corresponding to the low and intermediate FRET state (Fig. 6b) and a simplified 1D representation of the same was used to highlight the translocation behavior globally under different experimental conditions (Fig. 6c, d,e and f). In S-phase extracts, most 1D histograms showed a wide range of FRET efficiencies spanning the low and intermediate FRET states, demonstrating the oscillatory nature of translocating molecules (Fig. 6c). Since Ku is also sometimes actively removed from DNA, we observed several molecules with bias towards the high-FRET state in S-phase extract. Under conditions which prevent NHEJ complex removal, such as G1 phase extract or ΔCtIP S-phase extract, most molecules appear to be locked in the low FRET state (Fig. Fig. 6d, e), albeit some molecules do exhibit translocation. In S-phase extracts supplemented with ATMi, the translocation between low and intermediate FRET state was evident with the probability density of the majority of 1D histograms distributed between the two states (Fig. 6f).These results suggest that MRN mediated translocation of Ku is counteracted by the presence of an ‘elastic’ barrier which we propose is DNA-PKcs.

## Discussion

NHEJ is the dominant DSB repair pathway in all phases of cell cycle and its components such as Ku and DNA-PKcs rapidly bind DNA ends and prevent resection. For error-free repair to occur via HR, the removal of the NHEJ proteins is essential. Here, we elucidate the mechanism by which MRN and CtIP remove the NHEJ machinery from DNA ends to enable resection, the commitment step for homologous recombination. To this end, we developed of single-molecule assays that allowed real-time imaging of Ku removal and resection within a physiologically complex cell-free extract that recapitulates all major DSB repair pathways. Notably, we describe a previously unrecognized “translocase” activity of the MRN complex which evicts the NHEJ machinery from DNA ends and we delineated its regulation by CDK and ATM phosphorylation (Fig. 7).

**Figure 7:**
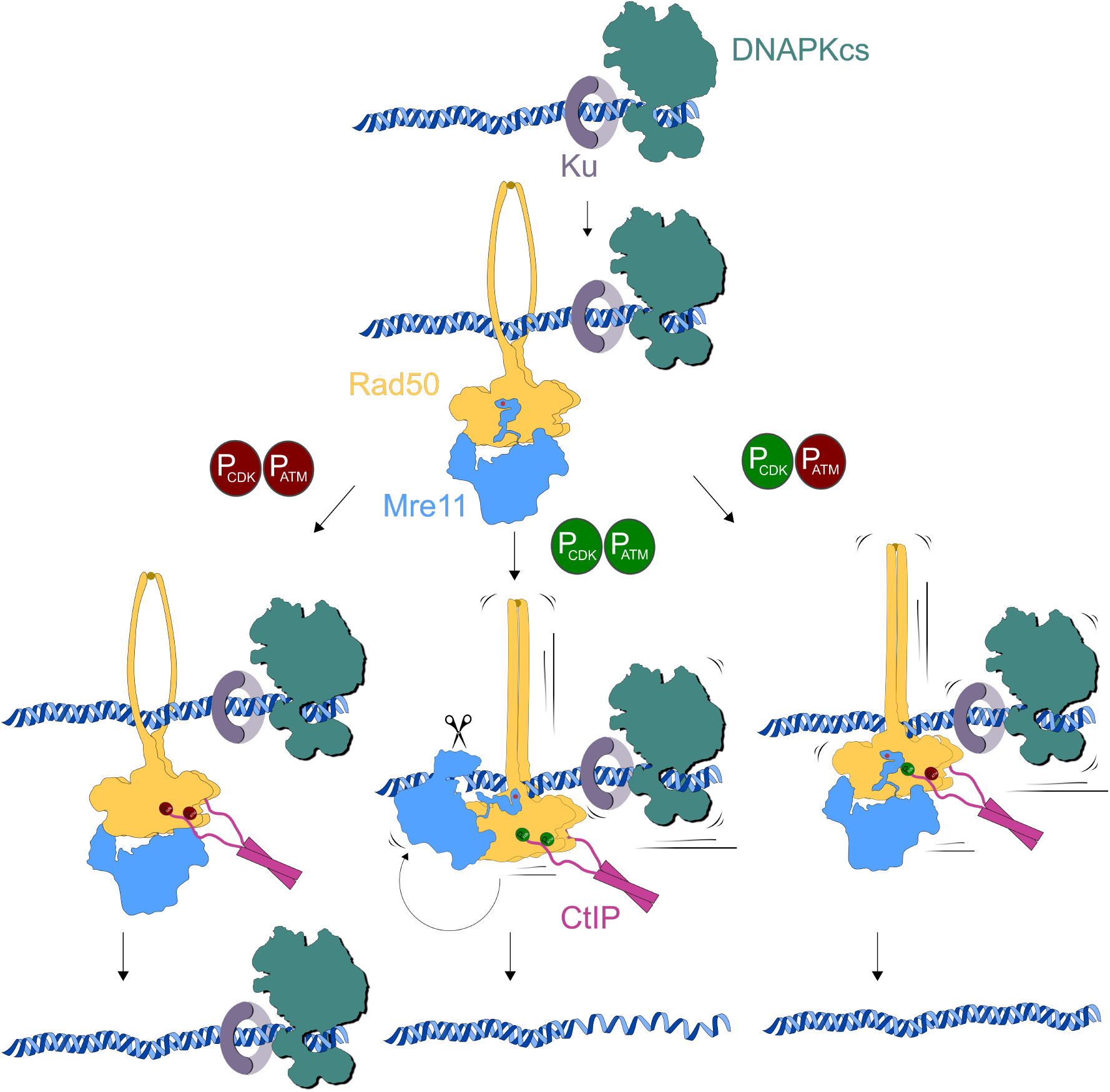
Model for mechanochemical removal of Ku by the MRN-pCtIP complex. Ku and DNA-PKcs are rapidly recruited to DSBs to protect the ends. In the S/G2 phase, CtIP is phosphorylated by CDK and subequently by ATM, which stimulates the endonuclease activity of Mre11. The cutting state involves repositioning of the Mre11 (Fig. 4b) and the collapse of Rad50 coiled coils around the DNA, the structural changes effecting 5^*′*^ end resection and Ku removal respectively. CtIP phosphorylation by CDK and ATM will culminate in the removal of Ku and resection of the 5^*′*^ strand (middle). Absence of both CDK and ATM phosphorylation results in an inactive conformation (left), devoid of structural rearrangement. In the case of ATM inhibition, Mre11 repositioning is hindered and therefore we observe no resection. However, the combined effect of Rad50 mediated translocation (loop extrusion) and constriction of the channel formed by collapsed coiled coils encircling the DNA can facilitate the removal of Ku from DNA ends in an ATP dependent manner (right).

We find that the dominant Ku removal pathway is mediated by the DNA resection machinery, which includes MRN and CtIP (Fig. 1e, f). Ku removal and end resection are highly coordinated with the removal of the NHEJ complex being necessary for resection (short-range) to proceed (Fig. 2a,b). Our experiments clearly indicate that resection towards the break proceeds via multiple nicks (as Ku is being pushed off), though 3^*′*^-5^*′*^ exonuclease activity by Mre11 [25] may also act in tandem. We see no evidence of dsDNA cleavage [33, 49].

MRN activity is tightly regulated by CtIP phosphorylation[24, 49, 22, 50]. CDK phosphorylation of CtIP is essential for eviction of the NHEJ complex, making it necessary but not sufficient for DNA resection (Fig. 3c). In contrast, ATM site mutants (CtIP3A) support Ku removal but block resection (Fig. 3d) These data, contrast to prior suggestions that DNA resection (by 3^*′*^-5^*′*^ Mre11 exonuclease activity) destabilizes the NHEJ complex on DNA ends [2, 51]. How do we reconcile the resection-independent removal of the NHEJ machinery with prior observations that Ku is stabilized in the presence of ATM inhibitor in cells [52]? In our experiments, fluorescently labeled NHEJ complexes were evicted when S-phase extract (containing unlabeled Ku) was added to the microchannel (Fig. 1a, d). Since ATM inhibition blocks resection, unlabeled Ku and other NHEJ components from the S-phase extract are expected to rapidly rebind to unbound DNA ends, preventing labeled Ku rebinding (Extended data Fig. 13). In conrast, in cells, Ku rebinding would result in a steady state level of Ku bound to DNA ends, giving the appearance that ATM inhibition stabilizes Ku on DNA after cell fixation and immunofluorescence analysis [35, 52].

Collectively, our data points towards Rad50 acting as a central hub receiving CDK- and ATM-mediated phosphorylation signals from CtIP to effect Ku removal and DNA end resection (Figure. 7). AlphaFold models show that the phosphorylated CDK (T847) and key ATM (T859) sites interact with distinct residues in the Rad50 N-terminal ATPase domain (R79 and K6 respectively) (Fig. 4b). Both sites are evolutionarily conserved and K6E is an embryonically lethal rad50s allele [43], a group of mutations that map to the N-terminal region of Rad50 which are defective in resection and clearance of Spo11 from breaks [53].

Rad50 belongs to the SMC family proteins and prototypical members like condensin, cohesin and SMC5/6 hydrolyze ATP to drive motor activity which is exploited for DNA loop extrusion [47, 46, 48]. In a similar manner, we speculate that NHEJ complex removal is achieved via ATP dependent Rad50 motor activity. Indeed, smFRET experiments with site-specifically labeled Ku clearly demonstrated that the NHEJ complex was actively translocated off DNA ends in S-phase extracts (Fig. 5c, e). The eviction of NHEJ complex was independent of resection as exemplified by efficient Ku removal in the presence of ATM inhibitor (Fig. 5d and 6f).

Assuming that Ku is translocated in a linear fashion devoid of significant orientational changes, the difference in the observed FRET efficiencies imply a translocation distance of ∼ 4 nm (12 bp). Cryo-electron microscopy structures of the core NHEJ factors[30] showed that within the long-range complex, a single copy of DNA-PKcs is positioned on the end threading a 11 bp stretch of dsDNA [30, 54]. Therefore, a 11 bp translocation of Ku towards the end is likely to destabilize DNA-PKcs, leading to its release from DNA. Without DNA-PKcs, Ku is no longer confined to DNA ends and is swiftly released, likely by MRN mediated translocation. DNA-PKcs is critical to limit Ku to a single-copy on DNA ends [37] and in its absence multiple Ku molecules load/unload rapidly from DNA (Extended data Fig. 14). Consistent with the tight association of DNA-PKcs with the DNA end and with the weak ATPase activity of MRN [55], we frequently observed ‘oscillatory’ motions of Ku prior to removal, spanning around 5 bp, which resulted from multiple attempts to translocate Ku towards the end followed by restoration to its original position upon unsuccessful removal of DNA-PKcs (Fig. 6a, Extended data Fig. 12). The oscillatory motions are a feature of active removal of the NHEJ complex from ends as conditions where Ku removal was inhibited, such as S-phase extracts depleted of CtIP/ATM inhibition/ATP*γ*S or G1 phase extracts (Fig. 6c, d, e), did not exhibit such motions and Ku remained stably locked in place (Fig. 5c).

An alternative model to translocation is that Rad50 extrudes a short segment of DNA, akin to other SMC family proteins. The progressive extrusion of DNA by Rad50, with the collapsed coiled-coils (encircling the DNA; Fig. 4a *right*) acting as a barrier for the NHEJ complexes, would result in a relative repositioning of the proteins towards the end before they are eventually ‘stripped off’ DNA ends. Interestingly, AFM imaging revealed that MRN when bound to DNA at internal regions made multiple contacts with DNA that resulted in loops of varying sizes [56].

In summary, we describe the mechanism by which the NHEJ machinery is evicted from from DNA ends and how this enables 5^*′*^ end resection. Our work raises a number of new questions regarding how the MRN complex carries out its distinct translocation and resection activities. Are motor proteins in addition to Rad50 necessary for efficient removal of the NHEJ machinery? How does phosphorylation of CtIP activate translocation and resection? Finally, how does the presence of the NHEJ synaptic complex impact the ability of MRN to disrupt the NHEJ machinery from DNA ends?

## Methods

### Egg extract preparation

Egg extracts were prepared using adult female *Xenopus laevis* (Nasco Cat LM00535). Frogs were cared for by the Center for Animal Resources and Comparative Medicine at Harvard Medical School (AAALAC accredited). Work performed for this study was in accordance with the rules and regulations set by AAALAC. The Institutional Animal Care and Use Committee (IACUC) of Harvard Medical School approved the work. G1 (High-speed supernatant; HSS) and S-phase egg extracts (Nucleoplasmic egg extract; NPE) were prepared as described [57].

### Assembly of flow cell

Two holes were drilled on either side of a cleaned glass slide (VWR, 48300-047) along the short axis. On either side of the pair of holes, strips of double-sided tape (Grace biolabs) were placed parallel to the short axis. A coverslip was then placed on the taped slide, with the biotin functionalized surface facing it to form a microchannel with the holes serving as inlet and outlet for buffer exchange. PE20 tubing was affixed to the inlet and outlet ports using epoxy. The outlet port was attached to gastight syringe (Hamilton) to draw solutions from PCR tubes containing protein/buffer/extract into which the inlet tube was placed.

### Fluorescent DNA substrates

Fluorescently labeled DNA substrates used in this study were generated by 1) PCR amplification using a 5^*′*^ biotinylated forward primer and a fluorescently labeled reverse primer or 2) by ligation of duplex containing a fluorescently labeled oligo to restriction enzyme digested DNA backbones (see Supplementary information for detailed protocol). pPS14 plasmid (modified pACEBAC vector) was used as the template for the PCR to obtain a 1.5 kb product. PCR reaction products were resolved on a 0.8 % agarose gel. The band corresponding to the product was excised and subjected to electroelution followed by ethanol precipitation. The DNA substrates were resuspended and stored in EB buffer (Qiagen).

### Fluorescent labeling of oligonucleotides

In a typical reaction, 2 µl of amino modified oligo (25 mg/ml) was labeled with 2 µl of NHS ester fluorophore derivatives (50 mg/ml), 10 % DMSO and 41 µl Sodium tetraborate buffer (100 mM pH 8.5). The reaction was incubated overnight at RT in the dark. The labeled oligos were ethanol precipitated and resuspended in 15 µl water and further subjected to UREA -PAGE in a 20% gel. The band corresponding to labeled oligos was excised and crushed by centrifugation through an Eppendorf containing a hole made by 18-gauge needle. 500 µl of EB buffer (Qiagen) was added to this crushed gel pieces and subjected to repeated freeze-thaw cycles. The tube was then rotated overnight at room temperature. The solution was collected after centrifugation and labelled oligo was recovered by ethanol precipitation.

### Protein purification

#### Halo-Ku80/Ku70

Ku80 coding sequence was amplified from plasmid described in earlier study [10] and subcloned into pACEBAC1 vector (Supplementary information). Ku70 was subcloned into pIDK vector. The two plasmids were then combined using crerecombinase to obtain a multigene plasmid pPS28. The plasmid was transformed into MAX Efficiency^™^ DH10Bac competent cells (ThermoFisher Scientific) and bacmid carrying the multigene plasmid was isolated by blue-white screening. Bacmid DNA was purified using a kit (ZR Bac DNA Miniprep Kit, Zymogen) according to manufacturer instructions. Sf-9 cells cultured in ESF 921 media (Expression Systems) were seeded in a 6 well plate (2 × 10^6^ cells). Sf-9 cells were transfected with HaloKu80/Ku70 Bacmid using Fugene Transfection reagent (Promega) according to manufacturer’s instruction and incubated for 72 h at 27^*°*^C. The viral particles were harvested by centrifugation at 500g for 10 min (P1). For amplification of the baculovirus, 0.5 ml of P1 was used to infect 25 ml of Sf-9 cells at 1×10^6^/ml for 96 hours at which point we observe ∼ 10 – 15 % cell death. The viral particles were isolated by centrifugation and further passed through a 0.22 µm syringe filter (P2) and stored in the dark at 4^*°*^C. For expression of the protein, 500 ml of Sf-9 cells at 2 × 10^6^ cells/ml were infected with the 0.5% (v/v) of the P2 for 72 h at 27^*°*^C. Cells were pelleted by centrifugation at 1000 g for 15 min at 4^*°*^C and finally resuspended in lysis buffer (50 mM Tris-HCl pH 7.5, 500 mM NaCl, 10 mM 2-Mercaptoethanol, 5 mM Imidazole, 10 % glycerol, 2 mM PMSF, cOmplete EDTA-free Protease Inhibitor Cocktail tablets). The cells were lysed by sonication with pulse cycle of 1s on and 6 s off, for a total on-time of 3 minutes at 40 % amplitude. The lysate was clarified by ultracentrifugation at 35000g for 1h and the supernatant was applied to Ni-NTA beads (Qiagen) (equilibrated in lysis buffer) at 4^*°*^C for 1h and placed on a rotating shaker. Beads were isolated by passing the lysate over a disposable column and non-specific proteins were removed by passing several volumes of wash buffer (50 mM Tris-HCl pH 7.5, 1M NaCl, 10 mM 2-Mercaptoethanol, 5 mM Imidazole, 10 % glycerol). Proteins were eluted using elution buffer (25 mM Tris-HCl pH 7.5, 500 mM NaCl, 10 mM 2-Mercaptoethanol, 300 mM Imidazole, 10 % glycerol). The eluates were pooled and diluted to bring down the salt concentration to 100 mM NaCl and then subjected to anion exchange chromatography using HiTrapQ-HP column (GE Healthcare). Fractions containing the Halo-Ku80/Ku70 complex were concentrated to around 0.4 ml using 10 kDa MWCO Amicon centrifugal filters (Millipore, UFC8010). The concentrated proteins were applied to a Superdex 200 Increase column (GE Healthcare) equilibrated in Ku SEC buffer containing 25 mM Tris-HCl pH 7.5, 300 mM NaCl, 1 mM DTT and 10 % glycerol. Fractions were analyzed by SDS PAGE and those corresponding to the Halo-Ku80/Ku70 complex were pooled, concentrated, snap-frozen in liquid nitrogen and stored at -80^*°*^C.

### CtIP

CtIP was expressed and purified from Sf-9 cells as described previously with modifications [16]. 500 ml of Sf-9 cells in suspension (2 m/ml) were infected with P2 baculoviral stocks as described earlier. After 60 h, the cell cultures were supplemented with 1 µM camptothecin (CPT; Sigma) and allowed to grow for 4h after which cells were pelleted by centrifugation. Cells were resuspended in 50 ml lysis buffer (50 mM Tris-HCl pH 8.0, 300 mM NaCl, 20 mM NaF, 20 mM Na_4_O_7_P_2_, 2 mM Na_3_VO_4_, 0.5% Triton X-100, cOmplete EDTA-free Protease Inhibitor Cocktail tablets and 10% glycerol) and lysed by sonication. The lysate was clarified by ultracentrifugation at 35000g for 1h and the supernatant was applied to ANTI-FLAG® M2 Affinity Gel (Sigma) (equilibrated in lysis buffer) and incubated in a rotating shaker at 4^*°*^C for 1h. The beads were transferred to a disposable column and washed extensively in wash buffer (25 mM Tris-HCl pH 8.0, 300 mM NaCl, 0.5 % Triton X-100 (Sigma), 10 mM β-Glycerophosphate and 10% glycerol). Bound protein was eluted with 5 column volumes of elution buffer (25 mM Tris-HCl pH 8.0, 150 mM NaCl, 100 µg/ml FLAG peptide (Sigma) and 10 % glycerol). Eluted protein was concentrated and snap frozen in liquid nitrogen and stored in -80^*°*^C until further use.

For In-vitro transcription-translation of CtIP and the various mutants used in this study, TnT SP6 High-Yield Wheat Germ Protein Expression System (Promega) was used according to manufacturer’s instructions.

### Fluorescent labeling of proteins

#### Halo-tag labeling

For labeling Halo-Ku70/80, 20 µM of the protein was incubated with 3-fold molar excess of Halo-JF549 (Promega) for 30 min at RT followed by overnight incubation at 4^*°*^C. The labeled proteins were separated from excess dye by gel filtration on a Superdex 200 increase column which was equilibrated with Ku SEC buffer. Resulting fractions were pooled, concentrated, and the degree of labeling was assessed prior to snap-freezing the proteins for storage at -80^*°*^C.

#### Ku80/Ku70-ybbr labeling

Ku80/Ku70-ybbr_338_ was purified as described above. For the labeling reaction, 20 µM of Ku80/Ku70-ybbr_338_ was supplemented with, 10 mM MgCl_2_, 10 µM Sfp synthase (plasmid obtained from Addgene (pET-Sfp, #159617) and purified as described [58]) and 40 µM CoA-Cy3 (prepared as described [58]). The reaction was incubated overnight at 4 ^*°*^C and the protein was separated from the free dye by gel filtration. The fraction containing proteins were pooled, concentrated and degree of labeling of assessed prior to snap-freezing the proteins for storage at -80^*°*^C.

#### Single-molecule Ku removal assay

20 µl of PBS containing BSA at 2 mg/ml (blocking buffer) was introduced into the flow cells to further block the surface and attenuate non-specific binding. Streptavidin (0.2 mg/ml in PBS; Sigma) was flown into the channel and incubated for 5 minutes at RT. The flow cell was washed with 60 µl of PBS to remove unbound Streptavidin. Biotinylated and fluorescently labeled DNA substrate was flown into the channel at a concentration ∼ 30-50 pM and incubated for 2 min at RT followed by washes with PBS to remove unbound DNA. The flow cell was placed firmly on the microscope stage for real-time visualization of all subsequent steps.

Ku depleted G1 egg extract was supplemented with Halo [JF549]-Ku (200 nM), ATP regeneration system (ARS; ATP (3 mM), phosphocreatine (15 mM) and creatine phosphokinase (0.01 mg/mL; Sigma)), NU7441 (100 µM; R&D Systems Inc). NU7441 was added to enhance the stability of the NHEJ factors on DNA ends. The extract was incubated at RT for 15 min and then flown into the microchannel. After 1 min of incubation the microchannel was flushed with ∼ 125 µl of egg lysis buffer (ELB; 10mM HEPES, pH 7.7, 50 mM KCl, 2.5mM MgCl_2_) to remove all traces of the extract. Approximately 70 % of DNA molecules contained NHEJ complex as visualized by the presence of fluorescently labeled Ku. Depending on the specific experiment, S-phase extracts were either immunodepleted using mock/Mre11/CtIP antibodies or supplemented with DMSO or inhibitors. ARS was added to the extract as described earlier and incubated at RT for 15 min. Immediately prior to flowing the extract into the microchannel, it was supplemented with Oxygen scavenging system (protocatechuic acid (PCA; 5 mM; Sigma); protocatechuate 3,4-dioxygenase (PCD; 0.1 mM; Sigma), methyl viologen (3 mM; Sigma) and ascorbic acid (3 mM; Sigma)).

#### Antibodies and immunodepletions

Rabbit polyclonal antibody against Ku80 used was described in an earlier study [10]. Polyclonal antibody was raised against full length recombinant *Xenopus* Mre11 protein by Pocono Rabbit Farm & Laboratory, Inc. Antibodies described above were used for western blot and immunodepletion.

All immunodepletions were carried out using the same procedure. 1x volume (packed) of Protein A Sepharose beads (GE Healthcare) was washed with PBS and incubated with 3x volume of the affinity purified rabbit polyclonal antibody (1 mg/ml) overnight at 4^*°*^C with constant mixing. The beads were washed in PBS buffer followed by ELB. 5x volume of egg extract was immunodepleted in three rounds with 1x volume of antibody coated beads for 1 h at 4^*°*^C with constant mixing. When using HSS, the extract was supplemented with nocadazole at 7.5 ng/µl. Immunodepleted extracts were aliquoted and flash-frozen in liquid nitrogen.

#### Pull-down assays

CtIP or mutant proteins were expressed in 15 µl WGE reactions (see above). Recombinantly purified xMRN (TwinStrept tag on Mre11) complex was added to this extract at a final concentration of 500 nM and incubated at RT for 30 min. 2.5 µl of MagStrep Strep-Tactin beads (IBA Lifesciences) was added to the reaction mixture and incubated for 1h at 4^*°*^C. Beads were washed in ELB (3x) and eluted in two rounds of 15 µl each using Strep-Tactin XT elution buffer (IBA lifesciences). 4 µl of eluate was separated by electrophoresis and detected by Western blotting using anti-CtIP or anti-Mre11 antibodies.

### Single-molecule image acquisition and analysis

#### Optical setup

Data presented in this study was collected from two microscopes. The optical layout of one of the microscopes has been previously described [59]. Here, imaging was carried out using a 100X objective and recorded using an EMCCD camera (Hamamatsu).

The second microscope is a home built objective type total internal reflection microscope. The optical setup was built around an Olympus IX-71 inverted microscope housing a 60x objective (Olympus PlanAPo 60X, 1.45 NA) and a filter cube fitted with a ZT532/640rpc dichroic and ZET405/488/561/640m emission filter (Chroma). Laser beams from a Coherent Sapphire 532-nm laser and a Coherent CUBE 641-nm laser were attenuated using a neutral density filter wheel (NDC-100S-4M; Thorlabs) and cleaned with a band pass filter (ZET 532/10X, ZET 640/10X; Chroma). The beam was expanded with a Galilean telescope constructed using an expanding and converging lens, following which they were combined using dichroic mirrors and expanded again with a pair of converging lenses and finally directed into the back port of the microscope. The laser beam was focused on the back-focal plane using a plano-convex lens mounted on a vertical translational mount which is in turn mounted on a lateral translational stage. TIRF angle was controlled by adjusting the vertical translation of the lens. Fluorescence emission exiting from the side port of the microscope was directed into a W-VIEW Gemini (Hamamatsu) image splitter which housed a ZT640rdc-UF2 dichroic (Chroma) and emission filter ET 595/50 (Chroma) and ET 685/50 (Chroma) for the green and red laser respectively, and finally focused on two halves of an sCMOS camera (ORCA-Quest, Hamamatsu). Illumination was controlled using Uniblitz VS14 shutters and driven by Uniblitz VMM-D3 shutter controller. TTL signals from the camera (master) were used to drive the shutters using a NI USB-6009 DAQ card (National instruments) in a synchronized manner. A motorized stage (Mad City Labs) was used to position the sample in the x-y plane and controlled by LabView software.

#### Imaging conditions

Fig. 1, Fig. 2a, e, Fig. 3, Fig. 4 and Fig. 5: Alternative stroboscopic illumination (excitation scheme: 4 frames green and 1 frame red) with exposure time of 200 ms and delay of 1s.

Fig. 2c: Simultaneous stroboscopic illumination with exposure time of 200 ms and delay of 1s

Fig. 6: Alternative continuous illumination (excitation scheme: 8 frames green and 1 frame red) with exposure time of 25 ms.

#### Image processing and data analysis

Python was used for all scripts related to image analysis and subsequent fluorescent signal processing in this study.

After image acquisition, data was stored in the form of `tiff` stacks. Images were processed using custom scripts to identify single molecules and extract the fluorescent intensity trajectories. Briefly, tiff stacks were loaded in the form of `Numpy` arrays using `Tiffile` (https://pypi.org/project/tifffile/). Image stacks were sliced according to excitation pattern to separate green and red channel emission signals and were corrected for stage drift using `chi2_shift` (https://github.com/keflavich/image_registration) where offset between reference and moving image are determined by DFT upsampling method. Typically, spot detection was performed on an ‘average’ image generated by averaging over the first 5 frames of the drift-corrected image stack. `chi2_shift` was used for image registration between the green and red channel ‘average’ images. A background image is first generated from the same stack by using a moving window for spatial averaging and applying gaussian filtering for denoising. The ‘average’ image was subjected to morphological processing, first with a gaussian filter to smoothen the image followed by ‘`dilation`’ operation to determine the locations of local maxima. Following this, statistical analysis was carried out by comparing the pixel values from the local maxima with the corresponding region in the background image to obtain preliminary peak positions of the single molecules. A box of pixels surrounding the peak location was clipped and intensity distribution was fitted to a two-dimensional Gaussian function by optimizing for least squares using the ‘Trust region reflective’ method. Sub-pixel coordinates of the single molecules were obtained along with the amplitude and standard deviation (σ) in the X and Y directions. Molecules which were separated by at least 2.5 σ and had a symmetric intensity distribution (σ_x_/ σ_y_ ∼ 1) were selected. Bright aggregates were discarded by a simple intensity threshold. Aperture photometry was performed on the image stack with background subtraction from an annular mask. Briefly, the image stacks are normalized for bit depth and the intensity values from a circular region of 5 pixels surrounding the Gaussian centroid are integrated to recover intensity for single molecule per frame. The local background is estimated by drawing an annulus with a width of 3 pixels around the circular area and sigma clipping (3σ, max iterations = 5) was performed to exclude outlier pixel values. Background corrected intensity values are used to obtain intensity traces shown in the manuscript.

Intensity traces derived from high-speed acquisition for Ku translocation (Fig. 6a and SI) were further processed by custom script for denoising by using modified version of the Chung and Kennedy filter. The ‘power’ term controlling the weight of the forward and backward predictors was adjusted to generate a smooth trendline in the FRET efficiency trace (Fig. 6a, middle panel).

#### Photobleaching analysis of Ku

G1 egg extracts depleted of endogenous Ku and supplemented with 200 nM Halo[JF549]-Ku was introduced into a microchannel containing the 1.5 kb DNA substrate tethered to the surface and incubated for 1 min. The channel was washed with 120 µl ELB and imaging was carried out in buffer containing oxygen scavenging system as described earlier (see methods). Step-detection was carried out using `quickpbsa` [60] where preliminary step identification was carried out using the Kalafut and Visscher algorithm followed by refinement using Bayesian approach. Half-step amplitudes that were supplied to the algorithm were estimated by custom scripts.

#### Survival probability curves

Survival probability curves were generated using the `lifelines` python package [61]. Step-detection was used to obtain the lifetimes of Ku or 5^*′*^ DNA from intensity traces. Data obtained from all experiments were right censored and a binary ‘censorship’ vector was passed as input for obtaining the final survival probability curve. Confidence interval (shaded area) for the estimated survival were computed at each data point using the Greenwood formula.

#### AlphaFold 3 multimer structure prediction

Sequences corresponding to human Mre11 (P49959), Rad50 (Q92878) and CtIP (Q99708) were obtained from UNIPROT database. A random 60 bp dsDNA was used for structure prediction (5^*′*^ AAAAATGCAAACTGCCGGCGAGCGTGCCGTT-TAGCCATAACATGGATACCATTGTGCCGG 3^*′*^). The sequences were submitted to the AlphaFold server (https://alphafoldserver.com) to generate models. To identify key interacting residues, contact map was obtained using https://mapiya.lcbio.pl/. Pymol was used for analysis and annotation of the structures generated from AlphaFold.

## Acknowledgment

This work was supported by Harvard Medical School Dean’s innovation grant (to J.J.L and J.C.W) and National Institutes of Health grant-R01CA272436 (to J.J.L). J.C.W. was supported by NIH grant HL098316. He is an American Cancer Society research professor and a member of the Howard Hughes Medical Institute. P.S and Y.W were supported by BCMP-Merck fellowship. We thank Dr. Tycho E.T. Mevissen and Dr. Geylani Can for assistance with Wheat Germ Extract and MRN pull down experiments. We thank members of the Loparo lab for useful discussions and critical comments on the paper.

