## Supplemental information for "The NHEJ machinery is translocated off DNA ends to enable resection"

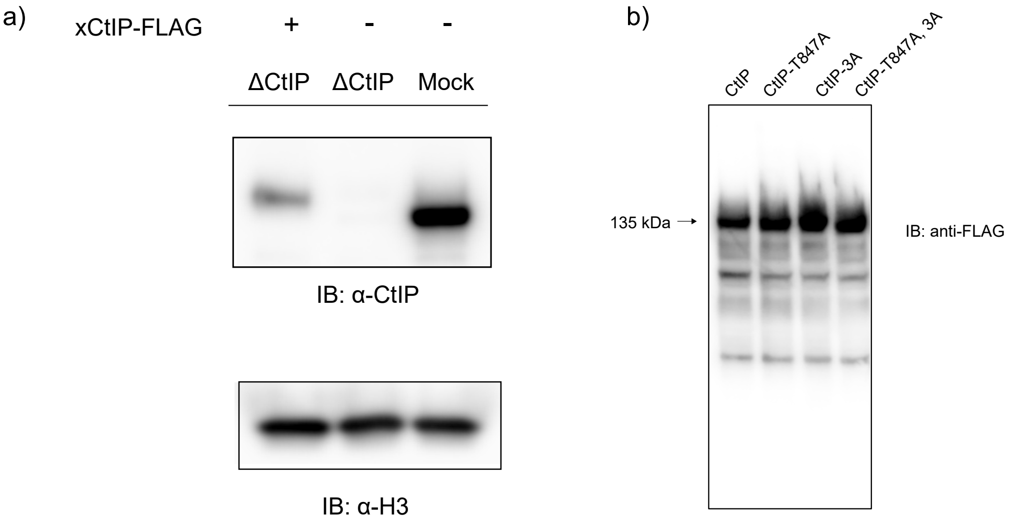


Figure 1: ***Western blot analysis of CtIP*** a) Western blot analysis of 1 µl Mock and and CtIP immunodepleted S-phase egg extracts(+/- Flag-CtIP) using anti-CtIP antibody. Histone H3 was used as loading control. b) Western blot analysis of Wheat Germ Extracts expressing CtIP-Flag or mutants using anti-FLAG antibody.

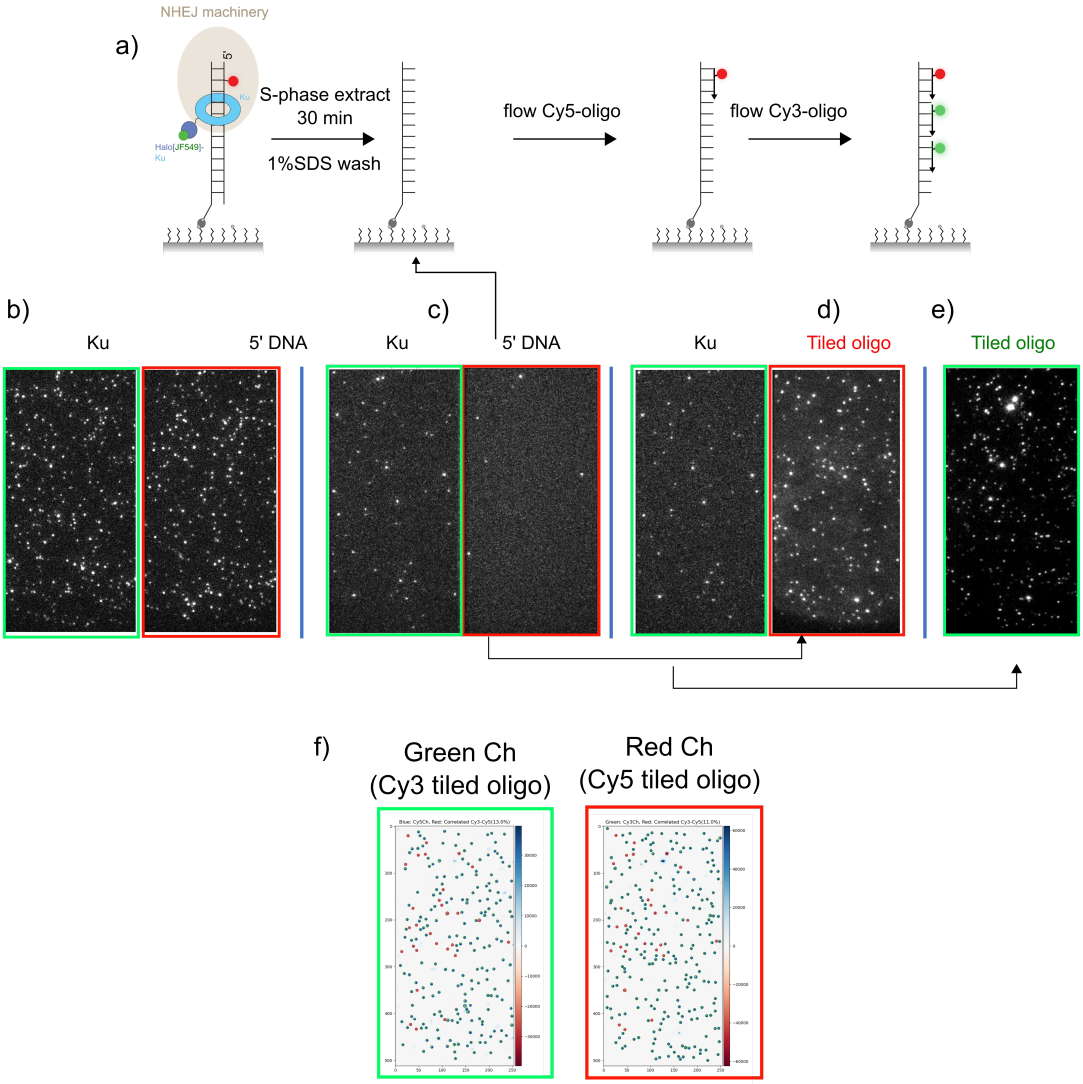


Figure 2: ***Resection in S-phase extract results in ssDNA overhang with intact 3′ end*** a) Schematic of experimental workflow. After incubating the DNA substrate with S-phase extract, the microchannel was washed with 1% SDS followed by copious buffer. Subsequently Cy3 or Cy5 labelled oligos were introduced in a sequential manner to hybridize to the ssDNA overhang generated as a result of resection. b) Images of Ku and DNA channels before and c) after incubation with S-phase extract. d) Fluorescence image after Cy5 labelled oligo complementary to the 3′ end was flowed into channel. e) Fluorescence image after Cy3 labelled oligos (3) were flowed into channel. Increase in spots shows binding of oligos to resected ssDNA. f) Colocalized Cy3 and Cy5 signal after D and E showing approximately 15 % molecules were bound by both sets of oligos.

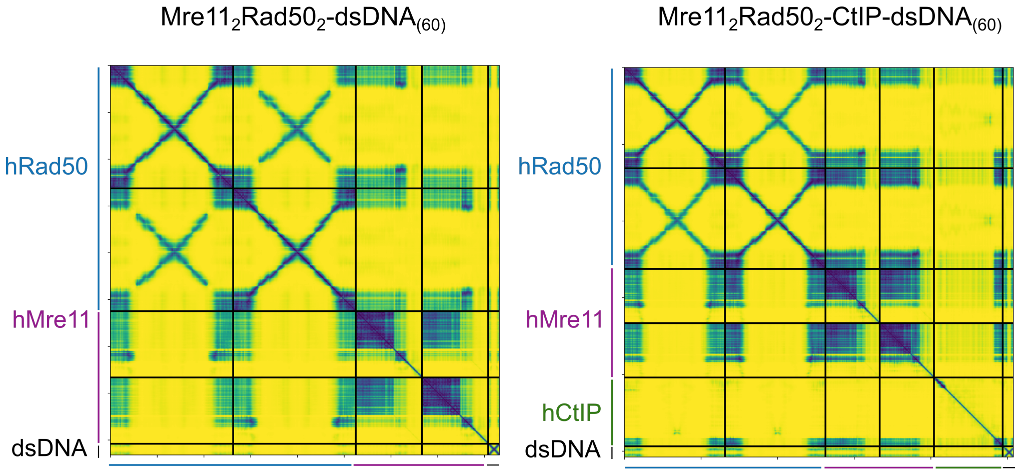


Figure 3: ***AlphaFold 3.0 (AF3) models of MR/MR-CtIP*** PAE plots of models generated in AF3 described in Fig. 4a. *left* MR with dsDNA molecule (60 bp) *right* MR-pCtIP with dsDNA molecule. The loss of interactions between N-terminal domain of Rad50 and Mre11 can be seen in the presence of phosphorylated CtIP owing to the repositioning of Mre11 (Fig. 4). Consequently, Mre11 interactions with dsDNA become apparent.

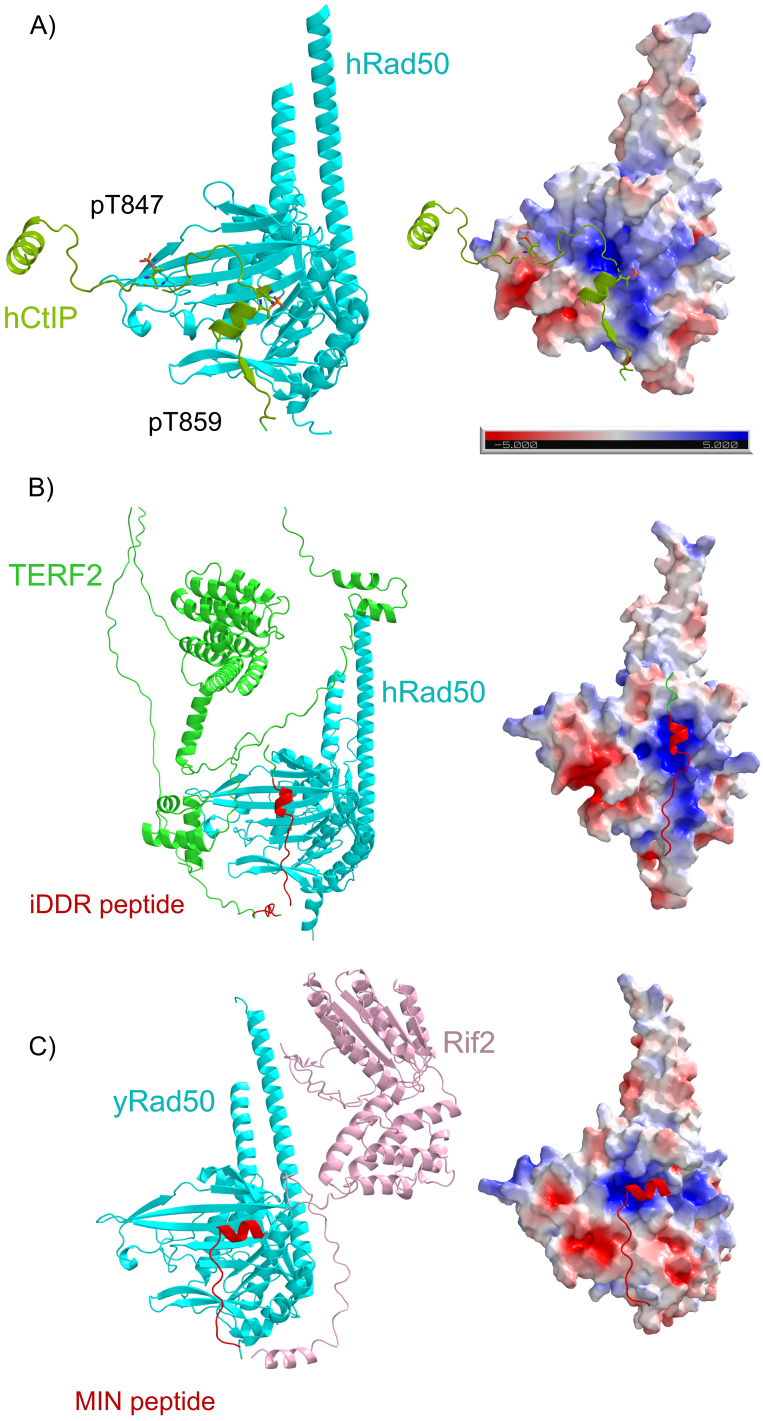


Figure 4: ***AF3 multimer models of Rad50 with interacting partners*** a) AF3 model of hRad50 with CtIP. Only residues with average PLDDT score > 70 are displayed. In addition, the N-terminal segment (1-140) is hidden for clarity. CDK (T847) and the key ATM (T859) site in the C-terminus of CtIP are labeled. The phosphorylated residues pack against a positively charged pocket in the N-terminal Rad50 β-sheets. b) AF3 model of hTERF2 and hRad50. The iDDR peptide (red) packs against the positive pocket in Rad50. c) AF3 model of yRif2 with yRad50 (MIN peptide; red).

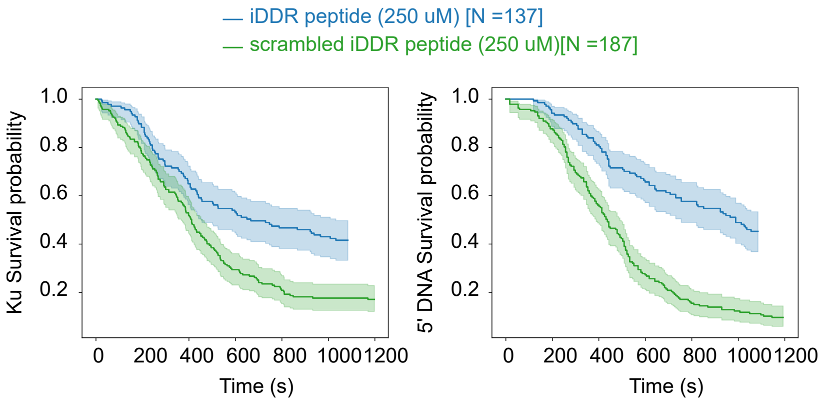


Figure 5: ***iDDR peptide inhibits Ku removal and resection*** Ku (*left*) and DNA (*right*) survival probability curves of S-phase extracts that were supplemented with 250 µM iDDR peptide or a peptide generated from the scrambled iDDR sequence

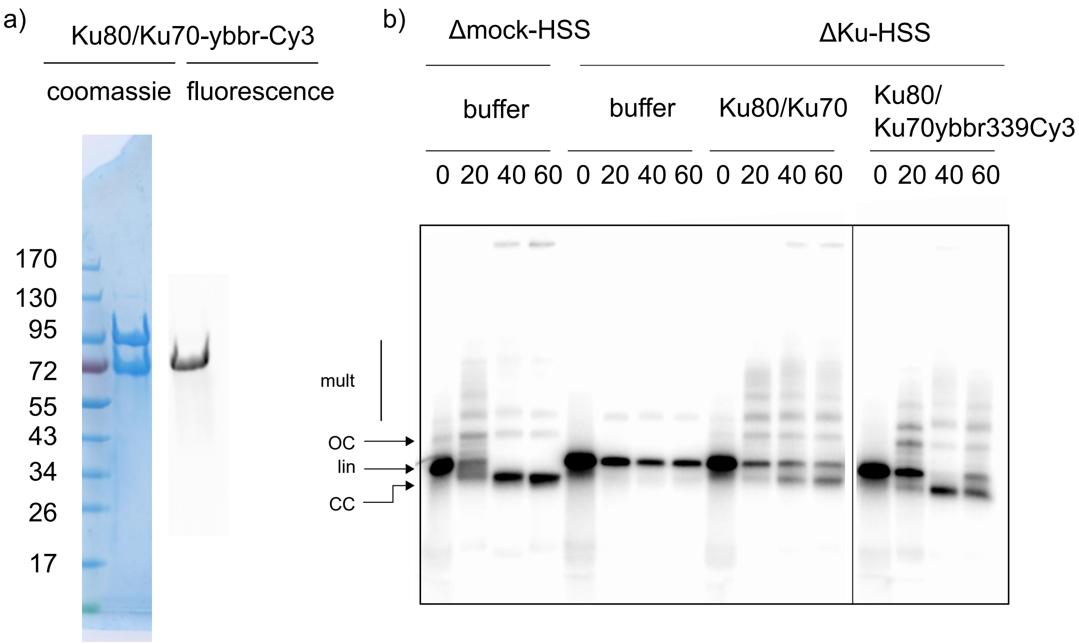


Figure 6: ***Purification of Ku80/K70-ybbr-Cy3 and assessment of activity*** a) Coomassie stained SDS-PAGE gel of purified xKu80/Ku70-ybbr-Cy3 and corresponding fluorescence image showing single band corresponding to fluorescent Ku70. b) A 3kb radiolabelled linear substrate was incubated in mock or Ku immunodepeleted G1 egg extracts supplemented with either buffer, Ku or Ku-ybbr as indicated (see methods). End-joining results in conversion of the linear (lin) substrate to supercoiled closed circular (Cc) form as observed from the difference in migration when resolved on a 0.8% agarose gel. Fluorescently labeled Ku is efficient in supporting end-joining.

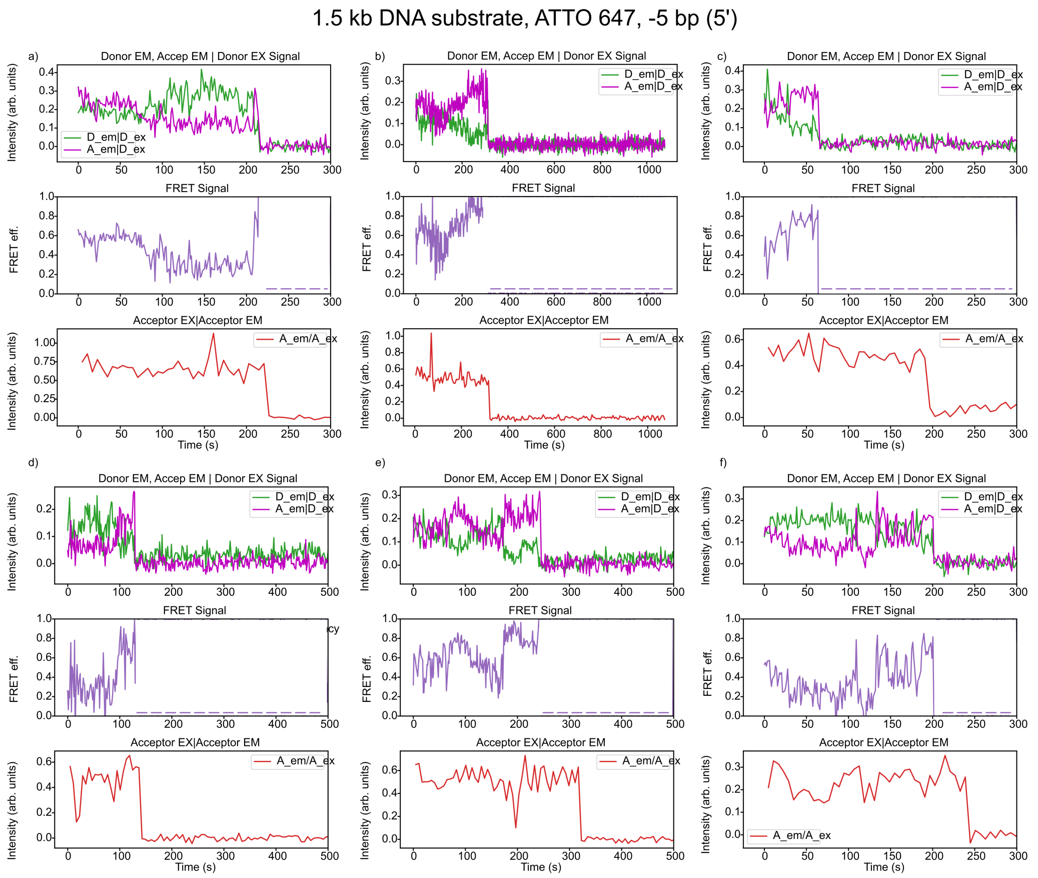


Figure 7: ***Single-molecule trajectories in S-phase extract*** a-f) Representative trajectories from experiment described in Fig. 5a and c.Trajectories show a transition from low-to high-FRET state prior to Ku removal, after which 5′ end is resected. The 1.5 kb DNA substrate was fluorescently labeled on the 5′ strand, 5bp from the end with ATTO 647.

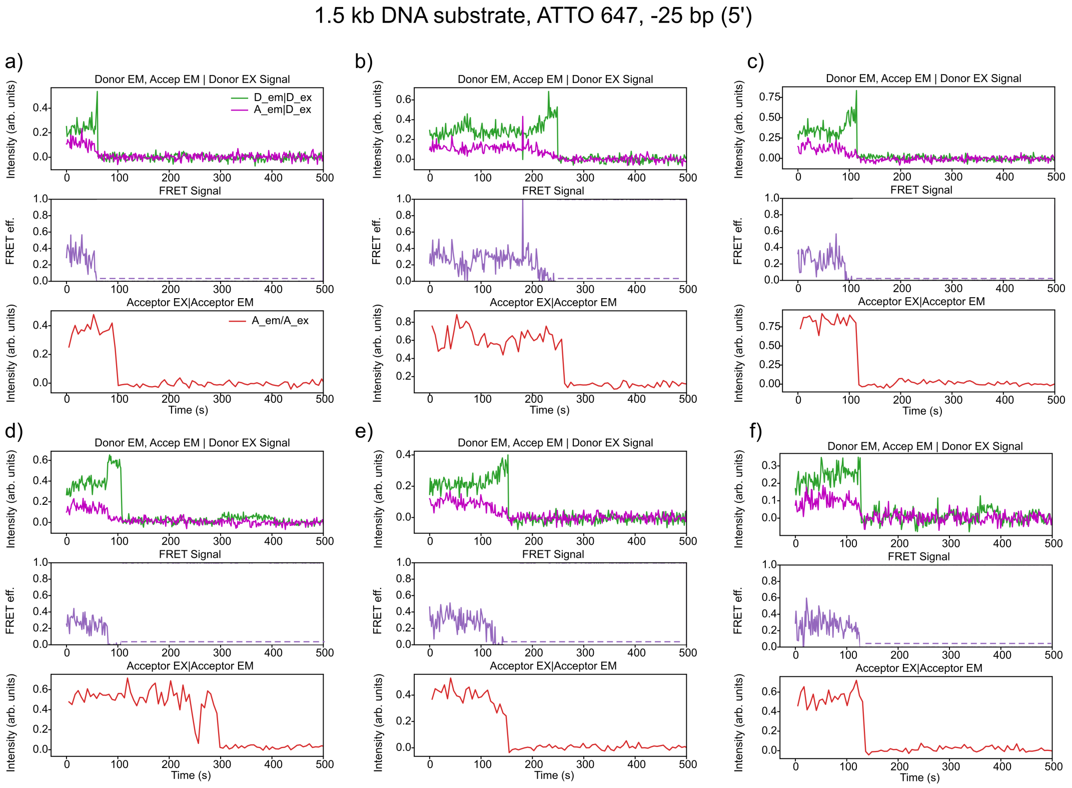


Figure 8: ***Single-molecule trajectories in S-phase extract*** a-f) Representative trajectories from experiment described in Fig. 5a except that the DNA substrate used has the fluorophore shifted to -25 bp from the -5 bp position. FRET trajectories show a decrease from mid- to low FRET state prior to Ku removal. 5′ end resection (*bottom panel*) occurs after Ku removal or on few occasions, simultaneously.

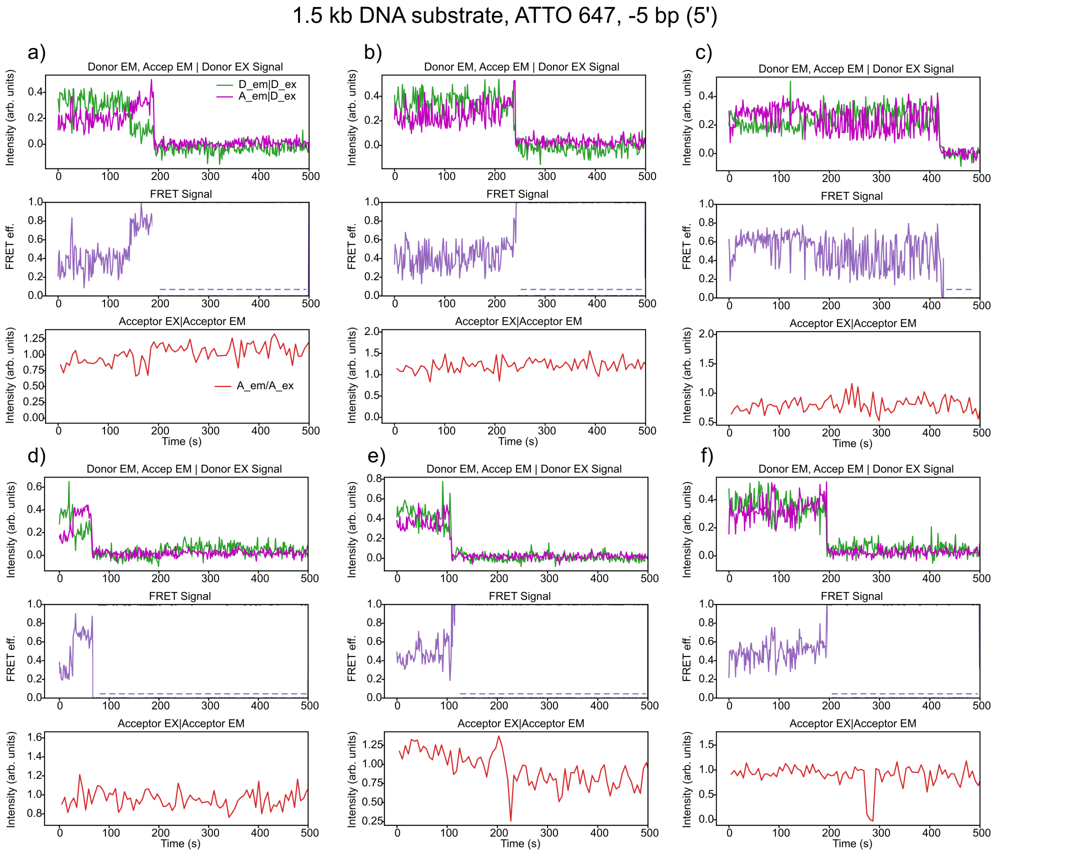


Figure 9: ***Single-molecule trajectories in S-phase extract supplemented with ATM inhibitor*** a-f) Representative trajectories from experiment described in Fig. 5d. Molecules show a transition from low-to high-FRET state prior to Ku removal and occurs in the absence of 5′ end resection (bottom panel). Enhanced fluctuations in the FRET efficiency trace reflect increased dynamics of Ku.

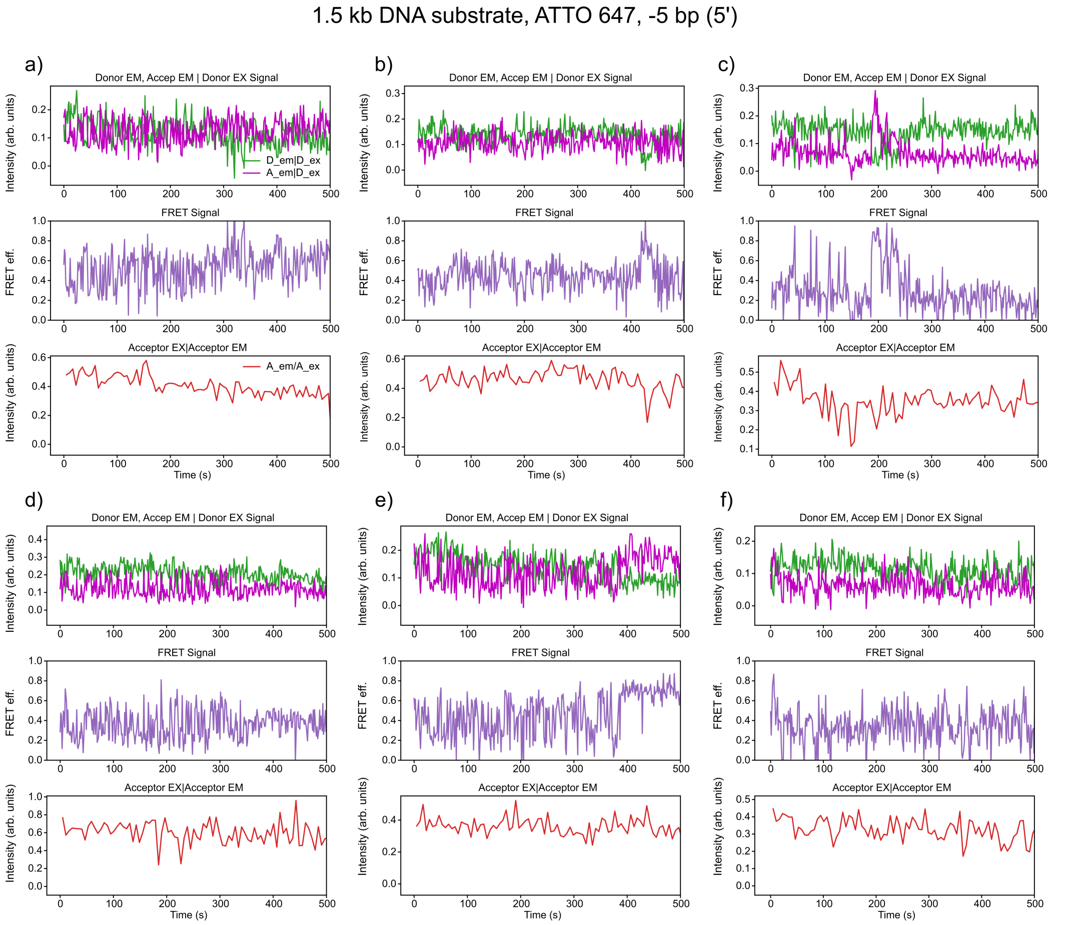


Figure 10: ***Single-molecule trajectories in $\Delta$CtIP S-phase extract supplemented with CtIP T847A*** a-f) Representative trajectories after supplementing CtIP depleted S-phase extract with CtIP T847A (see methods). Ku appears to be largely locked in the low-FRET state and exhibits no translocation highlighting that CtIP phosphorylation by CDK is indispensable for the process.

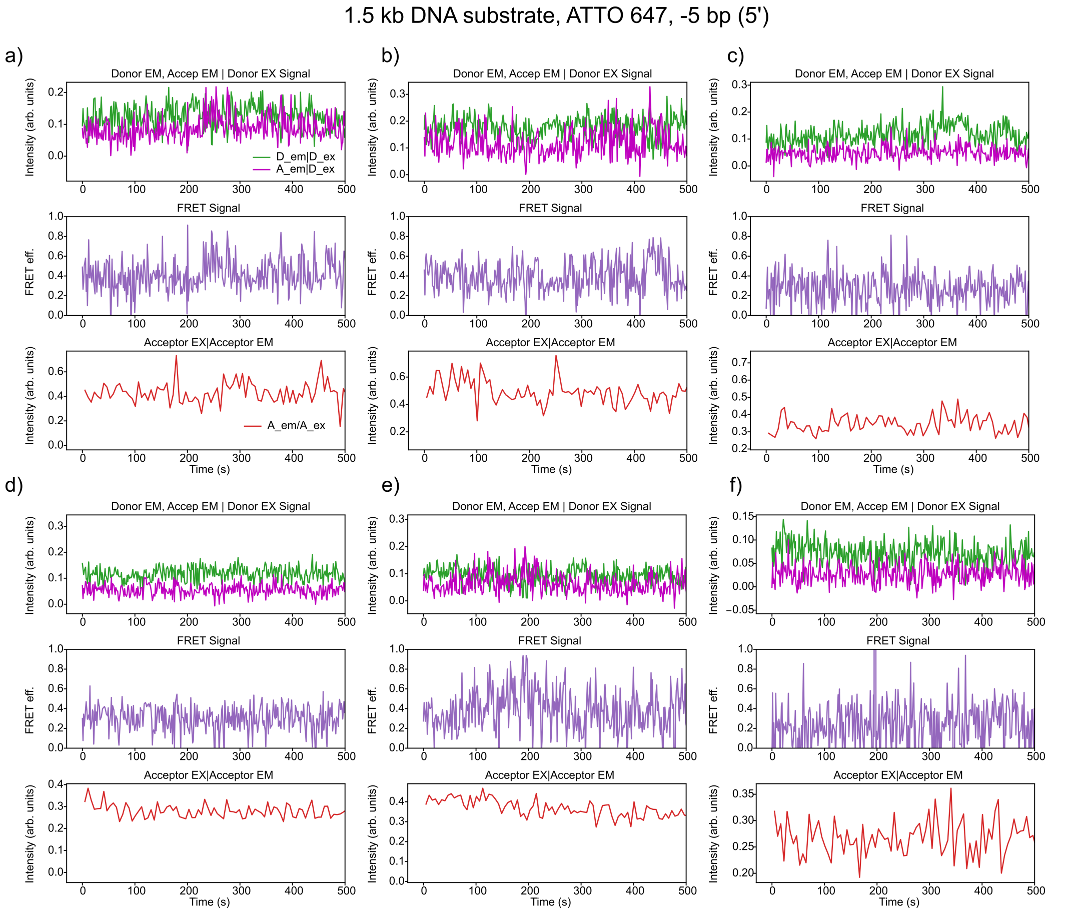


Figure 11: ***Single-molecule trajectories in S-phase extract supplemented with ATP$\gamma$S*** a-f) Representative trajectories after spiking S-phase extract with 3 mM ATP$\gamma$S. Ku appears to bind in the low FRET state and is not translocated off DNA ends, possibly due to inhibition of Rad50 ATPase activity.

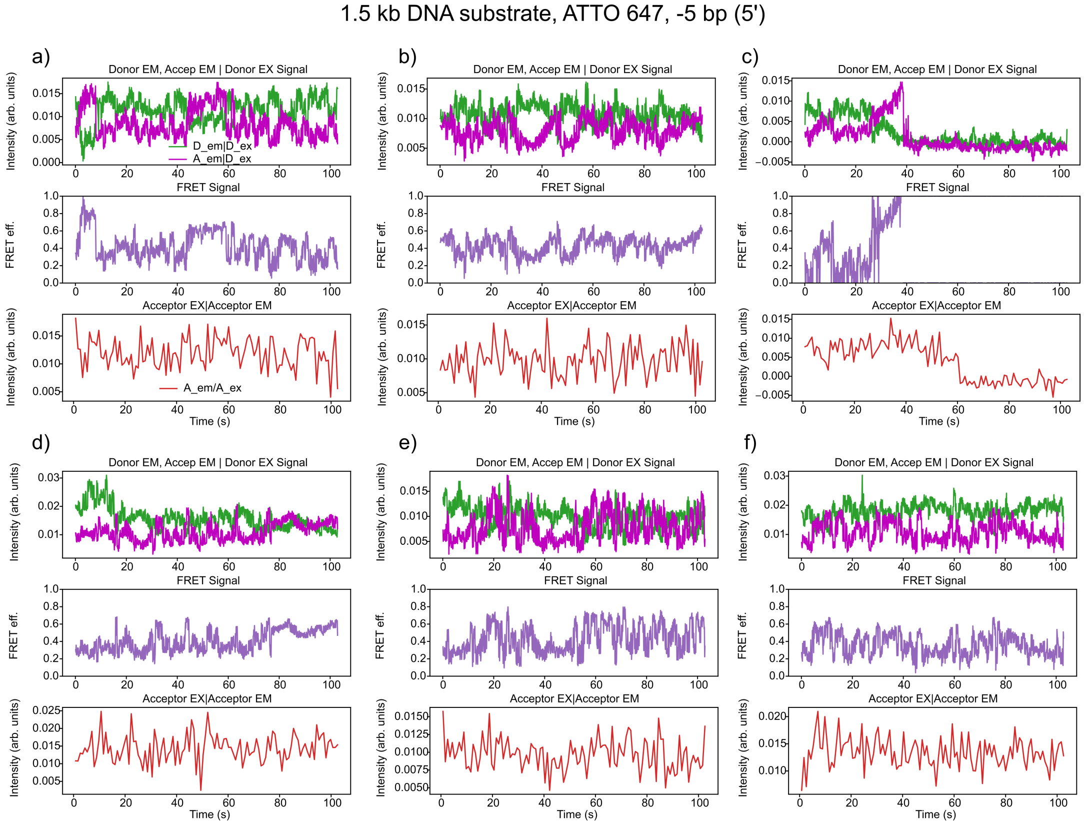


Figure 12: ***Single-molecule traces obtained from S-phase extract at high temporal resolution*** a-f) Representative trajectories which highlight the "oscillatory" motion of Ku molecules in S-phase extracts as described in Fig. 6a (see methods). Multiple oscillatory cycles are observed in each trace with varying periodicity. The anticorrelated donor and acceptor emission (*top panel*) intensity traces clearly point towards FRET between the two molecules.

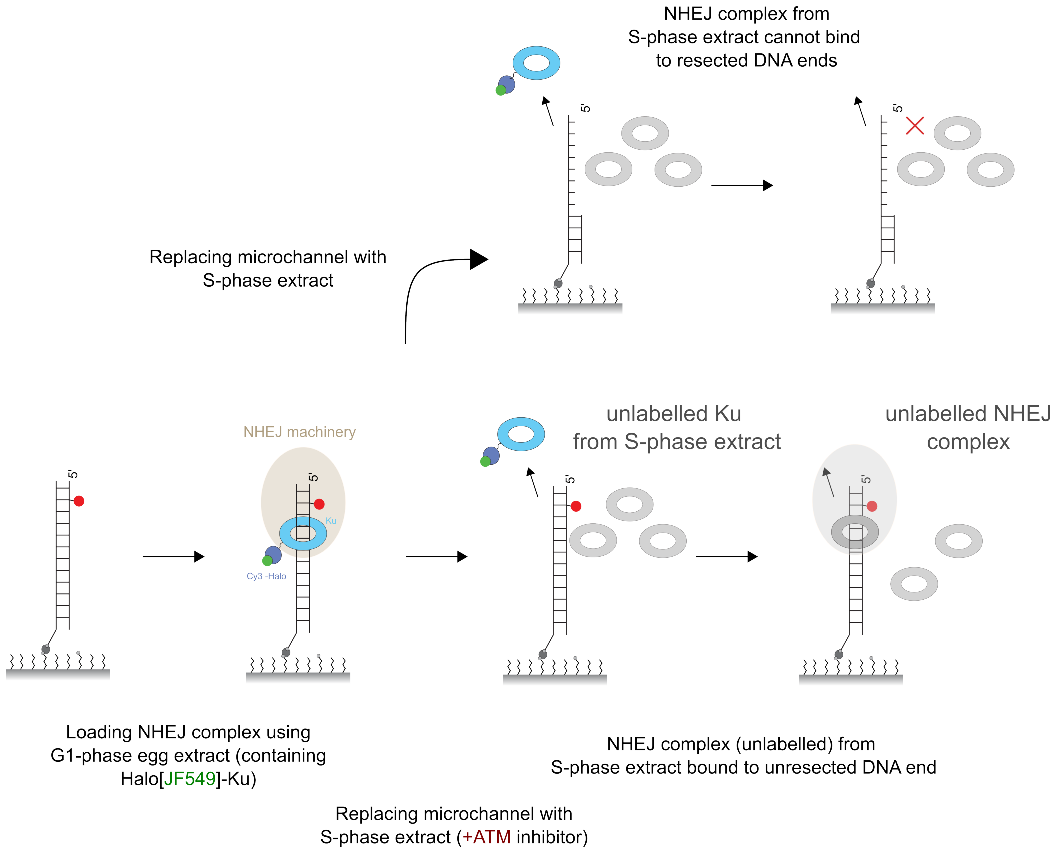


Figure 13: ***Apparent Ku stabilization in cellular experiments with ATM inhibitor*** In a typical single-molecule experiment, NHEJ complex is loaded on the DNA substrate using a G1 egg extract where endogenous Ku has been immunodepleted and supplemented with fluorescently labeled Ku. In the presence of S-phase extract, labeled Ku is removed and the DNA end is resected which suppresses teh loading of unlabeled Ku from the S-phase extract. However, in ATM inhibitor supplemented S-phase extract, resection is inhibited while Ku is removed efficiently. Consequently, Ku (in the S-phase extract; grey) can now rebind the DNA, undergoing multiple cycles of binding and removal.Therefore imaging Ku in fixed cells treated with ATM inhibitor (Britton et al., 2020) would show an apparent stabilization of Ku foci, since the dynamical nature of the process is imperceptible in these end-point experiments.

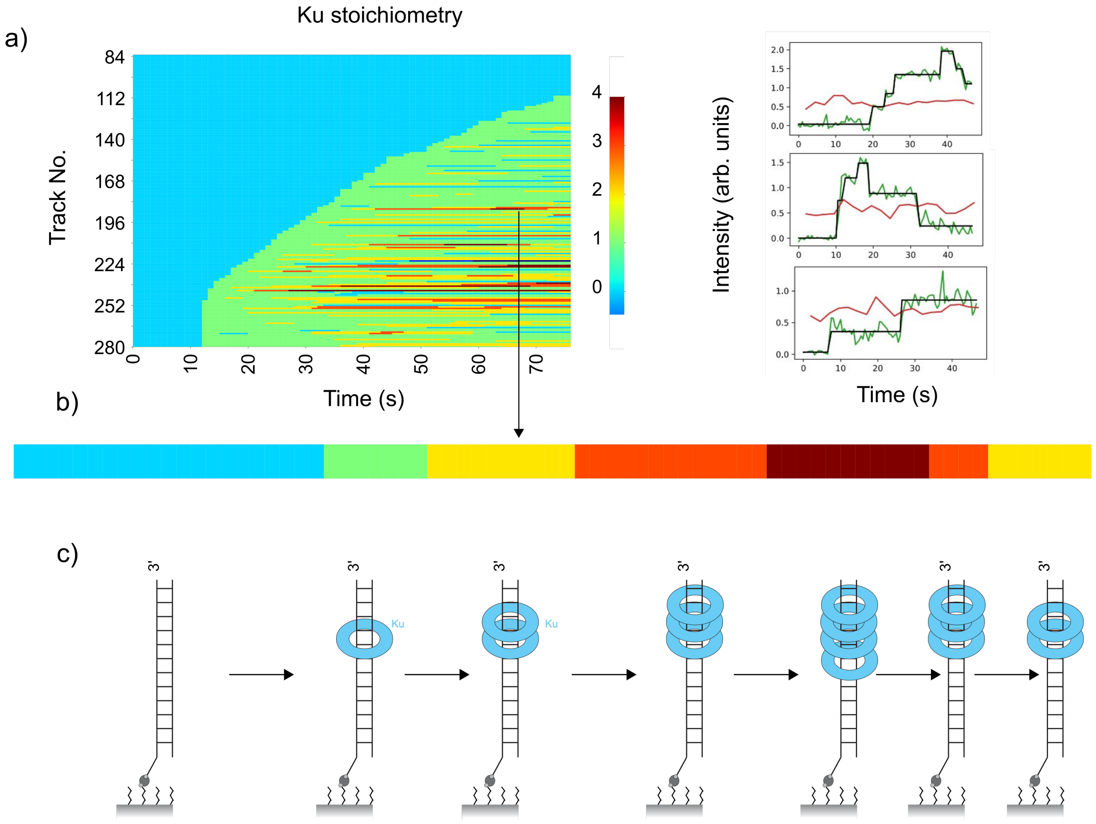


Figure 14: ***Multiple Ku molecules bind and rapidly slide along DNA*** a) Rastergram of 1 nM Halo[JF549]-Ku in ELB binding to 1.5 kb fluorescently labeled DNA substrate. The colours indicate the number of Ku molecules present. Ku binds and slides along the DNA facilitating the loading of another Ku molecule. In some instances, upto five molecules of Ku are threaded onto the DNA. Some representative raw traces are shown on the right. b and c) Example of one trace where four Ku molecules are loaded sequentially, following which we observe individual Ku molecules slide off the DNA end.
